# Loss of the intracellular enzyme QPCTL limits chemokine function and reshapes myeloid infiltration to augment tumor immunity

**DOI:** 10.1101/2022.01.25.477769

**Authors:** Rosa Barreira da Silva, Ricardo Leitão, Ximo Pechuan, Scott Werneke, Jason Oeh, Vincent Javinal, Yingyun Wang, Wilson Phung, Christine Everett, Jim Nonomiya, David Arnott, Cheng Lu, Yi-Chun Hsiao, James T. Koerber, Isidro Hotzel, James Ziai, Zora Modrusan, Thomas Pillow, Meron Roose-Girma, Jill M. Schartner, Mark Merchant, Sascha Rutz, Céline Eidenschenk, Ira Mellman, Matthew L. Albert

## Abstract

Tumor-associated macrophages are composed of distinct populations arising from monocytes or tissue macrophages, with a poorly understood link to disease pathogenesis. Here, we demonstrate that mouse monocyte migration was supported by glutaminyl-peptide cyclotransferase-like (QPCTL), an enzyme that mediates N-terminal modification of several subtrates, including the monocyte-chemoattractants CCL2 and CCL7, protecting them from proteolytic inactivation. Knockout of *Qpctl* disrupted monocyte homeostasis, attenuated tumor growth and reshaped myeloid cell infiltration, with loss of monocyte-derived populations with immunosuppressive and pro-angiogenic profiles. Antibody blockade of the receptor CSF1R, which more broadly eliminates tissue macrophages, reversed tumor growth inhibition in *Qpctl^−/−^* mice, and prevented lymphocyte infiltration. Modulation of QPCTL synergized with anti-PD-L1 to expand CD8^+^ T cells and limit tumor growth. QPCTL inhibition constitutes an effective approach for myeloid cell-targeted cancer immunotherapy.

## INTRODUCTION

Intratumoral accumulation of monocytes and macrophages has been associated with poor prognosis for cancer patients ^1, 2^. Within solid tumors, infiltrating monocytes give rise to a variety of macrophage populations, which may promote metastasis ^3^, angiogenesis ^4^ and resistance to immuno- and chemotherapy ^5–7^. Conversely, tumor-associated macrophages also support T cell responses, phagocytosis and tumor stroma remodeling, three mechanisms that promote anti-tumor immunity ^8–10^. Recent high-resolution profiling of myeloid cells in human tumors revealed an extensive heterogeneity of macrophage populations, which originate from infiltrating monocytes or tissue-resident macrophages ^11^. Although this heterogeneity suggests that different myeloid populations may be associated with tumor-promoting or anti-tumor activities, therapeutic strategies to reshape monocyte/macrophage populations in cancer remain elusive.

The various tumor-promoting functions of monocyte/macrophages in tumors have provided the rationale for the development of drugs targeting their survival and differentiation, such as m-CSF1/CSF1R neutralization ^12^. To date, these strategies demonstrated efficacy in the treatment of tenosynovial giant cell tumors, but have failed to demonstrate meaningful clinical benefit beyond this indication ^13, 14^ possibly due to a compensatory increase in the infiltration of immunosuppressive neutrophils ^15^, diminished availability of cytokines that support lymphocyte survival ^16^ and/or loss of beneficial CSF1R-expressing pro-inflammatory macrophages ^17^. To more selectively modulate the tumor monocyte/macrophage components, current approaches aim to impair monocyte migration, by targeting individual chemokine/chemokine receptors such as CCL2/CCR2 ^3, 18^. However, this strategy remains challenging due to the existence of multiple ligands and their pairing with several chemokine receptors ^19^, with several phase Ib/II clinical trials discontinued due to lack of pharmacodynamic effect or efficacy ^20, 21^. Strategies that can address this redundancy are needed to meaningfully test the role of monocyte-derived cells in tumor immunity, and to discern their functions from those of differentiated tissue macrophages.

Inhibition of the N-terminal X-proline dipeptidylpeptidase 4 (DPP4) protects multiple chemokines from inactivation, including CXCL10 and CCL11, resulting in enhanced anti-tumor immunity through T cells and eosinophils, respectively ^22, 23^. Given that the main monocyte chemoattractant proteins (MCPs), CCL2 and CCL7, possess the same N-terminal X-proline motif, we reasoned that similar pathways could be utilized to modulate their stability and thus limit monocyte accumulation into tumors.

## RESULTS

### QPCTL protects monocyte chemoattractants from DPP4-mediated truncation and inactivation *in vivo*

CCL2 and CCL7 are homeostatic chemokines detectable in mouse plasma at steady state (**Extended Data Fig. 1a**), and regulate monocyte homeostasis ^24, 25^. Both chemokines possess the N-terminal X-proline motif that can serve as a substrate for DPP4-mediated degradation (**Extended Data Fig. 1b**) but, contrasting to mice deficient in CCL2 and CCL7 ^25^, *Dpp4*^−/−^ mice had normal monocyte numbers in the blood, spleen and bone marrow (**Extended Data Fig. 1c**). Similarly, no monocyte phenotype was found in tumor bearing animals exposed to DPP4 inhibitor (DPP4i) **(Extended Data Fig. 1d)**. Previous reports proposed that cyclization of the N-terminal glutamine amino acid (Q) into pyroglutamic acid (pE), which results in a 17 Dalton reduction in molecular weight due to the loss of a -NH_3_ group, prevents DPP4- mediated truncation of the N-terminal glutamine-proline (QP) dipeptide of CCL2 and CCL7 ^26^ (**Extended Data Fig. 1e).** However, the relative abundance and role of these N-terminal chemokine-variants *in vivo* has not been assessed.

We developed immunoassays using newly generated antibodies specific for either the N-terminal cyclized (pE) or N-terminal truncated (ΔQP) forms of the chemokines (**Extended Data Fig. 1f–i**). Commercially available antibodies were used to establish a reference value for chemokine levels, referred to as ‘total– chemokine’. To parse the respective roles of the two known N-terminal cyclases: glutaminyl-peptide cyclotransferase (QPCT) and glutaminyl-peptide cyclotransferase-like (QPCTL) ^26, 27^, we generated *Qpct*^− /−^, *Qpctl*^−/−^ as well as *Qpct*^−/−^*Qpctl*^−/−^ double knockout mice. Applying these immunoassays, we established that the intracellular Golgi-localized QPCTL has a non-redundant role to protect CCL7 and CCL2 from N-terminal truncation (**Fig. 1a,b**). In contrast, the extracellular enzyme QPCT provided minimal contribution to chemokine modifications (**Fig. 1a,b**), despite being critical for extracellular glutaminyl-peptide cyclotransferase activity (**Extended Data Fig. 2a**). Loss of *Qpctl* reduced the amount of pE–CCL7, which is dominant in WT and *Qpct^−/−^* mice (**Fig. 1a**). We were not able to accurately measure pE–CCL2 at steady state (**Fig. 1b**). However, we detected pE–CCL2 in plasma from tumor-bearing mice, where total– CCL2 levels are higher, and identified a role for QPCTL in CCL2 N-terminal cyclization, as previously reported^27^ (**Extended Data Fig. 2b**). *Qpctl* deletion elevated total–CCL7 levels (**Fig. 1a and Extended Data Fig. 2c**), suggesting that CCL7 clearance and/or scavenging may be compromised, as previously described for DPP4-truncated CCL22 ^28^. Total–CCL2 levels were similar between the different genotypes (**Fig. 1b and Extended Data Fig. 2b**), suggesting that the regulation of CCL2 and CCL7 clearance may be different.

**Fig. 1.**
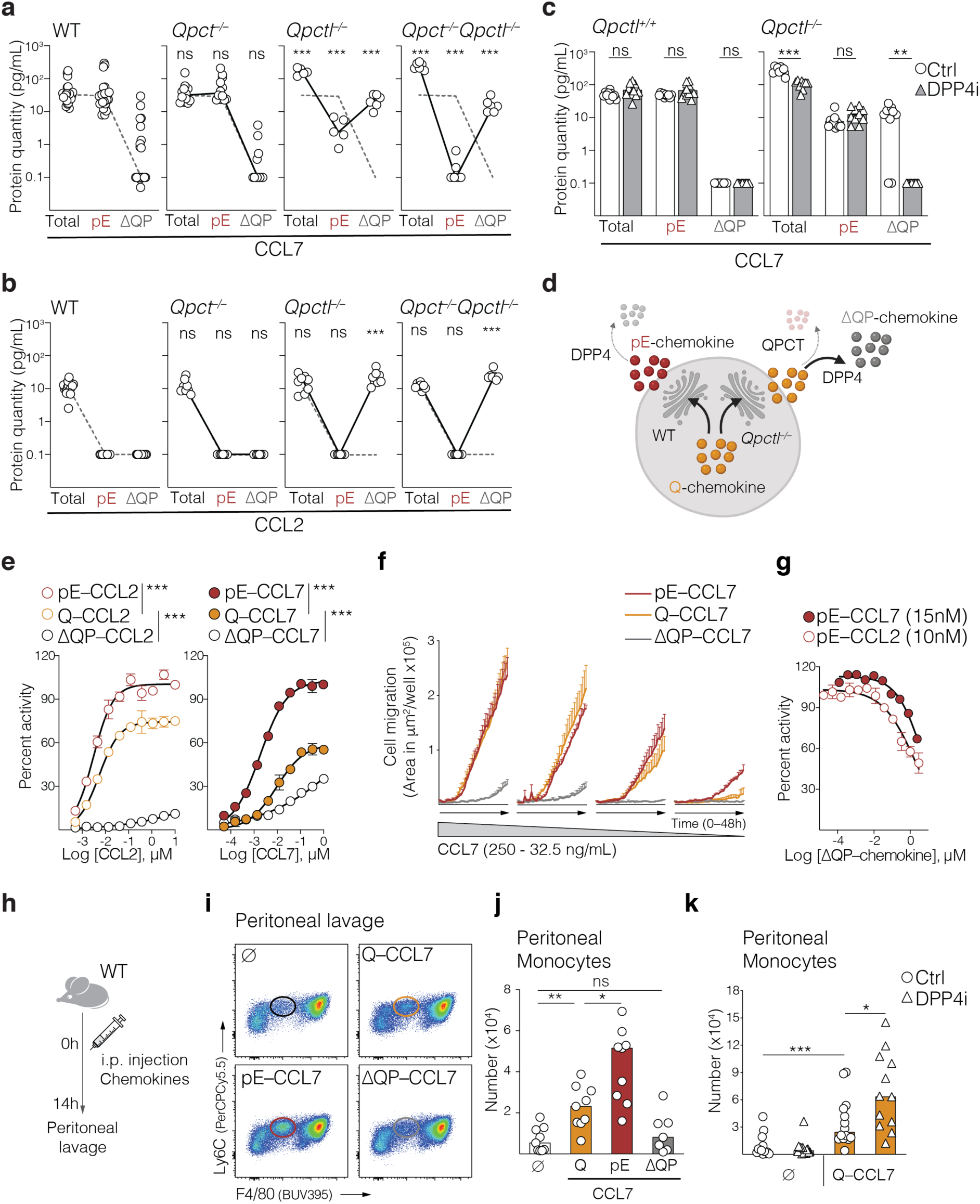
QPCTL protects monocyte chemoattractants from DPP4-mediated truncation and inactivation *in vivo*. **a**, CCL7 N-terminal post-translational modifications (PTMs) were quantified in the serum of naïve littermate *wild type* (WT, n = 30), *Qpct^−/−^* (n = 17), *Qpctl^−/−^* (n = 12) and *Qpct^−/−^Qpctl^−/−^* (n = 6) mice. Lines connect median values. Black dotted line represents median values from WT mice and statistics represent comparisons to WT values. **b**, CCL2 PTMs were quantified in the plasma of naïve littermate *wild type* (WT, n = 24), *Qpct^−/−^* (n = 8), *Qpctl^−/−^* (n = 8) and *Qpct^−/−^Qpctl^−/−^* (n = 8) mice. Lines connect median values. Black dotted line represents median values from WT mice and statistics represent comparisons to WT values. **c**, *Qpctl^+/+^* and *Qpctl^−/−^* littermate mice were treated with control (ctrl) chow or chow containing DPP4 inhibitor (DPP4i) sitagliptin. CCL7 PTMs were quantified in the serum, 4 weeks after treatment (n = 10 mice per treatment group). **d**, Schematic representation of *in vivo* N-terminal PTMs of monocyte chemoattractants. At steady state, the chemokine is cyclized at the Golgi by QPCTL, and secreted in its pE–form, resistant to truncation by the extracellular enzyme DPP4. In the absence of QPCTL (*Qpctl^−/−^*), the native chemokine in its Q–form is secreted. In the extracellular space, Q-chemokine is readily truncated by DPP4, rather than cyclized by the extracellular QPCT. **e**, The activity of chemokine forms in CCR2-expressing CHO-K1 cells was evaluated by measuring beta-arrestin recruitment. Stimulation was done with increasing concentrations of recombinant human (h)CCL2 or hCCL7 N-terminal modified forms. Line represents means and error bars SEMs (n = 2 technical replicates per condition). **f**, Transwell migration of differentiated human THP-1 cells to the indicated concentrations of hCCL7 forms was evaluated over time (48h) with the Incucyte technology. Line represents means and error bars SEM (n = 3 technical replicates per condition). **g**, Antagonist activity of ΔQP–hCCL2 and ΔQP–hCCL7 was evaluated in CCR2-expressing CHO-K1 cells stimulated with 10nM pE–CCL2 and 15nM pE–CCL7, respectively. Line represents means and error bars are SEMs (n = 2 (CCL7) or 6 (CCL2) technical replicates per condition). **h**,**i**,**j**, WT mice were challenged with an intraperitoneal injection of PBS (ø) or mouse (m)CCL7 chemokine forms. **i**, Representative flow cytometry plots (gated on live, singlets, CD11b^+^SiglecF^−^Ly6G^−^ and analyzed for indicated markers) are shown. Circles indicate monocytes. **j**, Bar graph depicting quantification of monocytes is shown (n = 9 or 8 (ΔQP) mice per group). **k**, WT mice treated with ctrl or DPP4i chow were challenged with an intraperitoneal injection of PBS (ø) or Q–CCL7. Quantification of peritoneal monocytes is shown (n = 13 (PBS WT), 14 (PBS DPP4i), 14 (CCL7 WT) and 12 (CCL7 DPP4i) mice per group). Bars are medians and symbols individual mice. Data shown is representative of one experiment (**b, e**-**g**) or pooled from 2 to 3 experiments (**a**,**c**,**j**,**k**). All experiments were repeated independently (⩾3 times). ns, not significant; \**p*⩽0.05; \*\**p*⩽0.01; \*\*\**p*⩽0.001. *p* values are from nonparametric Mann-Whitney test (**a**-**c**,**j**,**k**) or Two-way ANOVA test (**e**).

We examined the role of DPP4 in chemokine N-terminal truncation. Treatment of *Qpctl*^−/−^ mice with DPP4i, or crossing the *Qpctl*^−/−^ and *Dpp4*^−/−^ mouse strains, diminished CCL7 N-terminal truncation and decreased total-chemokine levels (**Fig. 1c and Extended Data Fig. 2d**). The concentration of pE–CCL7 remained low in *Qpctl*^−/−^ mice, regardless of DPP4 activity (**Fig. 1c and Extended Data Fig. 2d**), establishing that QPCT does not compensate for QPCTL loss. Moreover, this indicated that QPCT could not outcompete DPP4 in its modification of secreted chemokine. Thus, QPCTL defines, through cyclization, the N-terminal modification status of MCPs *in vivo*, protecting them from DPP4-mediated truncation (**Fig. 1d**).

To evaluate the functional consequences of N-terminal processing of MCPs we employed reporter assays for CCR2 signaling, which measure β-arrestin recruitment following receptor engagement. Both N-terminal pE– and the native Q– forms of the human chemokines triggered receptor signaling and β-arrestin recruitment (**Fig. 1e**). The pE–chemokines had the strongest receptor agonism, while N-terminal truncation resulted in loss of activity (**Fig. 1e**) and chemokine inactivation in migration assays **(Fig. 1f and Extended Data Fig. 2e),** extending prior observations ^26, 27^. Additionally, we observed a competitive inhibition of full-length chemokine-induced β-arrestin recruitment in the presence of truncated chemokines (**Fig. 1g**). Inhibition of chemokine-mediated AKT phosphorylation provided additional evidence for an antagonist activity of truncated chemokine (**Extended Data Fig. 2f**).

Using a model of intraperitoneal inflammation (**Fig. 1h and Extended Data Fig. 3a**), we observed that injection of recombinant mouse Q–CCL7 resulted in selective accumulation of monocytes in the peritoneal cavity, while truncated chemokine failed to do so (**Fig. 1 i,j and Extended Data Fig. 3b,c**). Recruitment of monocytes was higher after pE–CCL7 injection (**Fig. 1 i,j**), or upon inhibition or genetic loss of *Dpp4* (**Fig. 1k and Extended Data Fig. 3d**). Thus, QPCTL-mediated MCP cyclization was not required for monocyte migration *per se*, but was superior to induce intracellular signaling and important for maintaining MCP activity *in vivo* in the presence of DPP4.

### QPCTL function is essential for monocyte homeostasis

Consistent with the homeostatic function of MCPs ^24, 25^ and our findings that *Qpctl^−/−^* mice had elevated levels of truncated, non-functional MCPs, *Qpctl^−/−^* mice also had fewer circulating and splenic monocytes (**Fig. 2a,b and Extended Data Fig. 4a,b**). This phenotype was associated with an increase in CD115^hi^ bone marrow monocytes (**Fig. 2c and Extended Data Fig. 4c)**, similar to the phenotype reported in mice lacking *Ccr2, Ccl2* or *Ccl7* expression ^24, 25^. Consistent with the dominant *in vivo* role of QPCTL in MCP modification, this phenotype was also observed in *Qpct*^−/−^*Qpctl*^−/−^ double knockout, but not in *Qpct^−/−^* mice (**Fig. 2d–f**). QPCTL appeared to play a specific role in monocyte hematopoiesis, as *Qpctl^−/−^* mice presented normal abundance and distribution of liver Kupffer cells, lung alveolar macrophages and brain microglia (**Extended Data Fig. 5a)**, which are established from fetal macrophages and monocytes, in a CCR2-independent manner ^29^. Additionally, lymphocyte and granulocyte populations were also unaffected (**Extended Data Fig. 5b**). Splenic patrolling monocytes were reduced (**Extended Data Fig. 5c**), likely a result of their development being partially dependent on classic bone marrow monocytes ^30^. When analyzing reciprocal bone marrow chimeras, we determined that lack of *Qpctl* expression in either the donor bone marrow (*Qpctl^−/−^*→WT) or in recipient stromal cells (WT→*Qpctl^−/−^*) acted to suppress monocyte egress to the blood and spleen (**Extended Data Fig. 5d**). Both reciprocal chimeras presented elevated levels of ΔQP–CCL7, when compared to WT controls (**Extended Data Figure 5e**), suggesting that monocyte homeostasis is dependent on chemokines secreted from both stromal and hematopoietic compartments. WT and *Qpctl^−/−^* monocytes co-transferred into a WT recipient migrated similarly to the peritoneal cavity (**Extended Data Fig. 5f**), indicating that *Qpctl^−/−^* monocytes are phenotypically normal with respect to CCL7-induced mobilization.

**Fig. 2.**
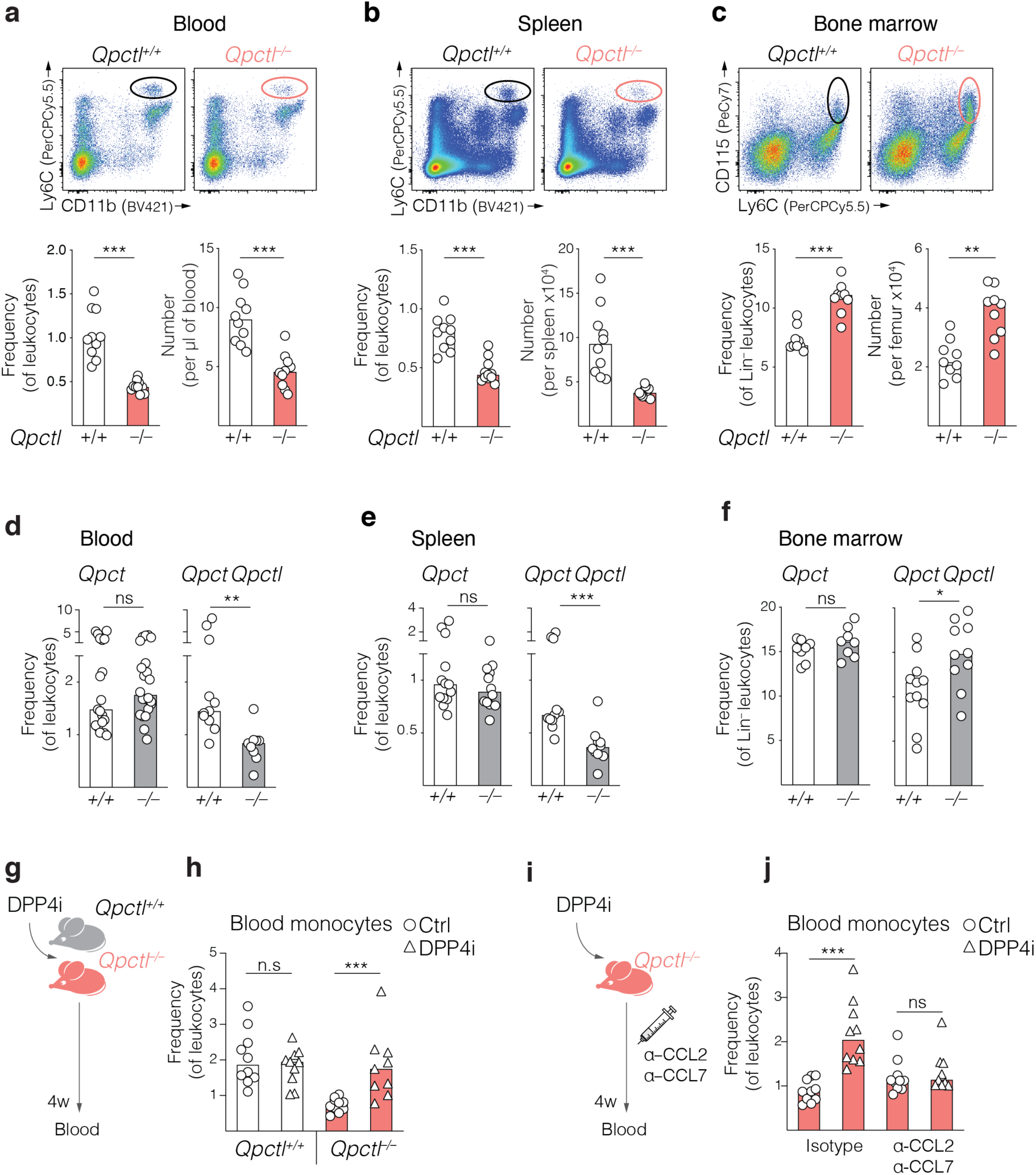
QPCTL function is essential for monocyte homeostasis. **a**,**b**,**c,** Analysis of Ly6C^+^ monocytes in **a**, blood, **b**, spleens and **c**, bone marrow of naive *Qpctl^−/−^* and WT littermates (*Qpctl^+/+^*). Representative flow cytometry plots (gated on live, singlets and analyzed for indicated markers) are shown. Circles indicate monocytes. Quantification of monocyte frequency and number is shown in bar graphs (**a**, n = 10 mice; **b**, frequency n = 10 or 11 (*Qpctl^−/−^*) mice; number n = 10 or 9 (*Qpctl^−/−^*) mice; **c**, n = 9 mice per group). **d**,**e**,**f**, Analysis of Ly6C^+^ monocyte frequency in **d**, blood, **e**, spleens and **f**, bone marrow of *Qpct^−/−^*, *Qpct^−/−^ Qpctl^−/−^* double mutant mice and respective WT littermates (**d**, left graph n = 19 or 22 (*Qpctl^−/−^*) mice; right graph n = 11 or 10 (*Qpct^−/−^Qpctl^−/−^*) mice; **e**, left graph n = 13 or 12 (*Qpct^−/−^*) mice; right graph n = 11 or 10 (*Qpct^−/−^Qpctl^−/−^*) mice; **f**, left graph n = 8; right graph n = 11 or 10 (*Qpct^−/−^Qpctl^−/−^*) mice per group). **g**,**h**, *Qpctl^−/−^* and WT littermate mice were treated with ctrl chow or chow containing DPP4i for 4 weeks. **h**, Frequency of Ly6C^+^ blood monocytes is shown (n = 10 or 9 (*Qpctl^−/−^* DPP4i) mice per group). **i**,**j**, *Qpctl^−/−^* mice treated with ctrl of DPP4i chow were injected with anti-CCL2 plus anti-CCL7 neutralizing antibodies, or with isotype controls. **j,** Quantification of monocytes in the blood, after 4 weeks of treatment is shown (n = 10 mice per group). Bars are medians and symbols individual mice. Data shown is pooled from 2-3 experiments. All experiments were repeated independently (⩾3 times). ns, not significant. \**p*⩽0.05; \*\**p*⩽0.01; \*\*\**p*⩽0.001. P values are from nonparametric Mann-Whitney test.

We hypothesized that if QPCTL facilitated monocyte mobilization by preventing chemokine inactivation via N-terminal truncation, then inhibition of DPP4 in *Qpctl^−/−^* mice would restore monocyte homeostasis. Indeed, inhibition or genetic loss of *Dpp4* increased monocyte frequency in blood and spleen of *Qpctl*^−/−^ mice (**Fig. 2g,h and Extended Data Fig. 5g,h**). Conversely, treatment with CCL2 and CCL7 neutralizing antibodies prevented DPP4i-mediated rescue of monocyte mobilization in *Qpctl*^−/−^ mice (**Fig. 2i,j**). This data demonstrates that QPCTL-mediated N-terminal modification of MCPs is critical for the development of monocyte niches, and thereby sustains monocyte homeostasis *in vivo*.

### Modulation of QPCTL selectively inhibits monocyte infiltration and tumor growth

Monocytes and monocyte-derived macrophages accumulate in solid tumors following chemoattractants ^3^ and constitute the main immune infiltrate in many cancers ^31^. To assess if QPCTL regulates monocyte migration to tumors, we selected the breast cancer EO771 and lung carcinoma LL/2 mouse tumor models, which expressed *Ccl2*, *Ccl7* and *Qpctl* **(Extended Data Fig. 6a)**, and presented a high frequency of infiltrating monocytes (**Extended Data Fig. 6b,c)**. We generated *Qpctl-*deficient variants of both models using CRISPR/Cas9. Bulk transfection of the cell lines resulted in >95% of *Qpctl* mutation and, as expected, *Qpctl^−/−^* cells were defective in pE–CCL7 production (**Extended Data Fig. 6d**), yet presented similar kinetics of growth *in vitro*, when compared with parental or WT control cells transfected with non-targeting Cas9 RNPs (**Extended Data Fig. 6e**).

We inoculated WT mice with WT tumor cells or *Qpctl*^−/−^ mice with *Qpctl*^−/−^ tumor cells and monitored tumor growth over time. Remarkably, loss of *Qpctl* expression attenuated the growth of both models (**Fig. 3a–c**). We further observed fewer tumor-infiltrating Ly6C^+^ monocytic cells in *Qpctl*^−/−^ groups (**Fig. 3d,e and Extended Data Fig. 7a**). There was no significant impact on the number of differentiated macrophages (identified by high expression of F4/80 and lack of Ly6C expression) or other immune cells (**Fig.3e and Extended Data Fig. 7a**).

**Fig. 3.**
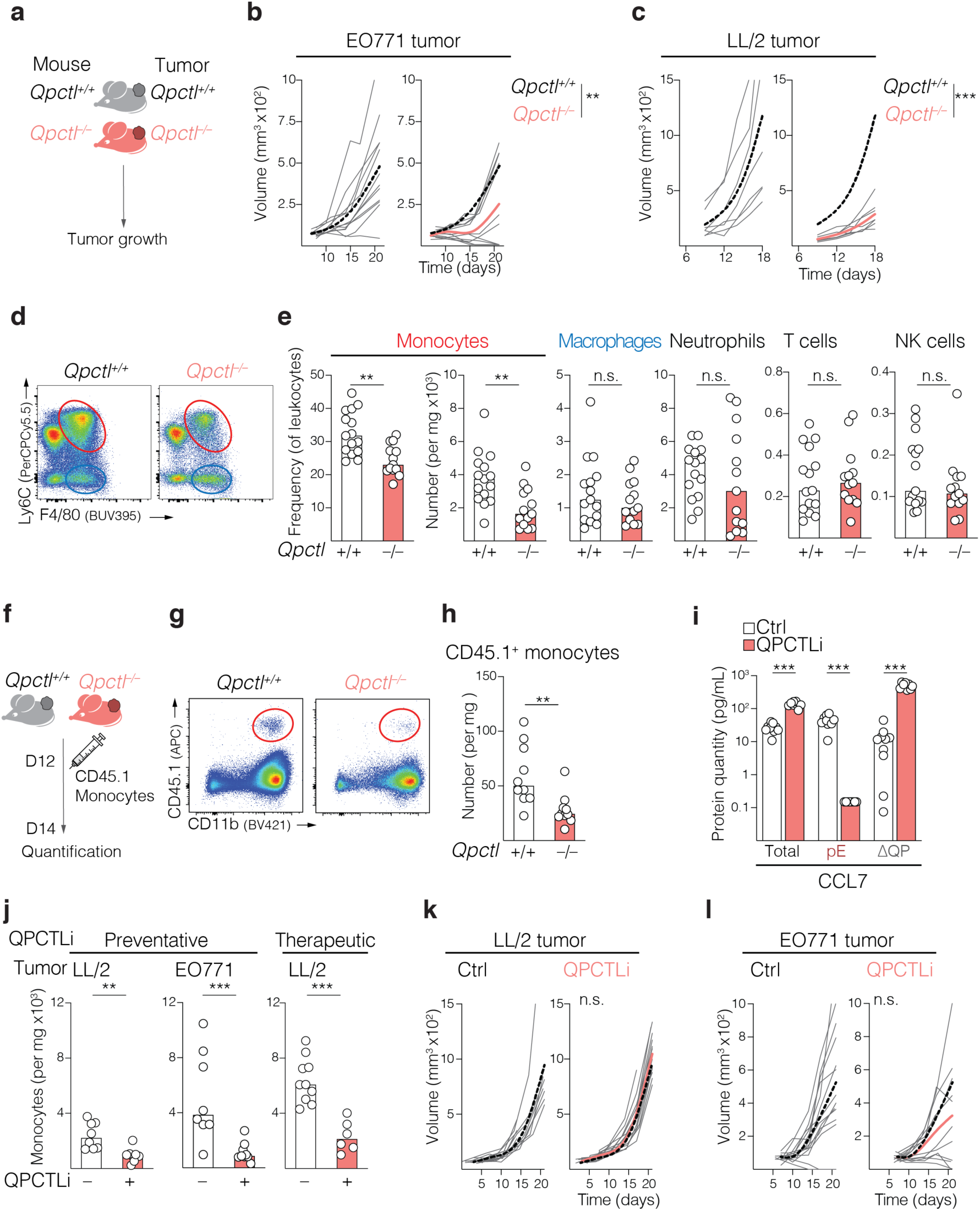
Modulation of QPCTL selectively inhibits monocyte infiltration and tumor growth. **a**,**b**,**c**, *Qpctl^−/−^* and *Qpctl^+/+^* littermate mice were inoculated with *Qpctl^−/−^* or *Qpctl^+/+^* tumors, respectively. **a**, Schematic representation of the experimental protocol. **b**, EO771 growth was measured over time (n = 12 mice per group). **c**, LL/2 growth was measured over time (n = 7 mice per group). Gray lines represent individual mice. Overlay of fitting spline curves for each group is shown. Black dotted lines represent spline curves from *Qpctl^+/+^* groups. **d**,**e**, LL/2 tumors were excised 14 days after tumor inoculation and analyzed by flow cytometry. **d**, Representative flow cytometry plots identifying Ly6C^+^ monocytic cells (red circle) and F4/80^hi^ macrophages (blue circle) among tumor infiltrating leukocytes. **e**, Frequency of tumor infiltrating Ly6C^+^ monocytes and number of Ly6C^+^ monocytes, macrophages, neutrophils, T cells and NK cells is depicted, n = 15 or 13 (*Qpctl^−/−^*) mice per group. **f**,**g,h,** *Qpctl^−/−^* and *Qpctl^+/+^* littermate mice were inoculated respectively with *Qpctl^−/−^* or *Qpctl^+/+^* LL/2 tumors. CD45.1^+^ WT monocytes were transferred into tumor bearing mice, 12 days post tumor inoculation. Tumors were analyzed 2 days after monocyte transfer. **f**, Schematic representation of the experimental protocol. **g**, Representative flow cytometry plots identifying transferred monocytes (red circle) among leukocytes. **h**, Quantification of transferred monocytes is depicted (n = 10 mice per group). **i**, CCL7 PTMs were quantified in the plasma of WT mice treated with ctrl vehicle or QPCTL inhibitor (QPCTLi) for 4 days (n = 10 or 8 (QPCTLi) mice per group). **j**, WT mice received ctrl or QPCTLi for 4 days before LL/2 or EO771 tumor inoculation (preventative) or at day 7 after LL/2 inoculation (therapeutic). On day 14 after tumor cell inoculation, tumors were excised and the number of tumor-infiltrating Ly6C^+^ monocytic cells determined by flow cytometry (n = 8 (LL/2 preventative); n = 8 or 9 (EO771 QPCTLi), n = 10 or 6 (LL/2 therapeutic QPCTLi) mice per group). **k,l**, WT mice received ctrl or QPCTLi starting 1 day before LL/2 or EO771 tumor inoculation. Tumor growth was measured overtime. n = 15 (LL/2 ctrl); n = 14 (LL/2 QPCTLi) n= 14 (EO771 Ctrl), n = 12 (EO771 QPCTLi). Gray lines represent individual mice. Overlay of fitting spline curves for each group is shown. Black dotted lines represent spline curves from the ctrl groups. Bars are medians and symbols individual mice. Data shown is representative of one experiment (**c**,**i**,**j,k,l**) or pooled from 2 experiments (**b**,**e,h**). All experiments were repeated independently (⩾2 times). ns, not significant, \**p*⩽0.05, \*\**p*⩽0.01, \*\*\**p*⩽0.001. P values are from nonparametric Mann-Whitney test (**e,h,i**) or Two-way ANOVA test (**b**,**c,k,l**).

As *Qpctl^−/−^* mice have fewer circulating monocytes (**Fig. 2a**), we wanted to disambiguate these developmental defects from the putative role of QPCTL in the inflammatory trafficking of monocytes into tumors. Transferred WT monocytes migrated less to *Qpctl^−/−^* tumors, demonstrating that loss of *Qpctl* impaired inflammatory mobilization of monocytes (**Fig. 3f-h**).

We next tested whether pharmacological inhibition of QPCTL could also limit monocyte migration to tumors. Glutaminyl cyclase inhibitors are being developed for the treatment of Alzheimer’s disease, to prevent the generation of pathogenic pyroglutamate-modified proteins^32, 33^. We employed a QPCTL inhibitor (QPCTLi) (Compound 11, WO2014140279), which prevented N-terminal CCL7 cyclisation when administered to mice (**Fig. 3i**), confirming inhibition of the enzyme *in vivo*. QPCTLi reduced Ly6C^+^ monocytic cell accumulation in both LL/2 and EO771 tumors (**Fig. 3j**), a result observed when administered either preventatively, before tumor cell injection, or therapeutically, when tumors were established. QPCTLi also reduced the number of splenic monocytes (**Extended Data Fig. 7c**), in agreement with the proposed role of QPCTL in monocyte homeostasis. In LL/2 tumors QPCTLi also reduced the number of differentiated macrophages (**Extended Data Fig. 7d**). While this result was expected given that monocytes differentiate into macrophages within the tumor, genetic deletion of *Qpctl* did not impair macrophage numbers in this model and we did not observe reduced macrophage numbers in EO771 tumors. The differences between genetic deletion and pharmacological inhibition of *Qpctl* in the subcutaneous LL/2 model may be due to phenotypic differences in the host mouse strain used (in-house *Qpctl^−/−^* mice and *Qpctl^+/+^* littermate controls versus commercially available C57BL/6 mice), which could account for differences in the kinetics and/or differentiation potential of monocytes into tumor macrophages or could reflect differences in the relative contribution of tissue-resident macrophages in mice with constitutive *Qpctl* deficiency versus acute QPCTL inhibition. Additionally, administration of QPCTLi did not impact LL/2 growth, while we observed a partial inhibition on EO771 tumors (**Fig. 3k,l**). While many factors could account for the weaker effect of the inhibitor, it is likely that acute QPCTL inhibition did not fully recapitulate, in particular early on, the phenotype of *Qpctl*^−/−^, thus explaining the observed weaker effect on tumor growth. The differences in macrophage numbers observed between *Qpctl^−/−^* and inhibitor-treated mice in the LL/2 model further support this hypothesis and may translate into different tumor growth kinetics.

This data indicates that QPCTL modulation preferentially impairs accumulation of recently-differentiated monocyte-derived cells, while sparing some populations of tissue-macrophages potentially insensitive to QPCTL activity.

### Anti-CSF1R treatment reverses the beneficial effect of *Qpctl* loss on tumor growth

Strategies to limit tumor monocyte/macrophage infiltration include administration of CSF1R-targeting molecules, which are currently in clinical development ^12, 17^. To understand how CSF1R targeting compares to *Qpctl loss*, we analyzed EO771 and LL/2 tumors growing in WT mice treated with anti- CSF1R antibody. Anti-CSF1R treatment reduced the number of Ly6C^+^ monocytic cells and eliminated F4/80^hi^ macrophages from the tumors (**Fig. 4a,b**), as described. The number of tumor-infiltrating lymphocytes, including T and NK cells, were also reduced (**Fig. 4b and Extended Data Fig. 7e,f**), suggesting that CSF1R-expressing myeloid populations were important to sustain lymphocyte infiltration. Anti-CSF1R treatment did not attenuate tumor growth, supporting previously published data ^15^ and highlighting the distinct mechanism of action achieved following *Qpctl* loss (**Fig. 4c,d**). In addition, anti-CSF1R treatment reversed EO771 and LL/2 tumor growth inhibition observed in *Qpctl*^−/−^ groups **(Fig. 4c,d)**. We therefore hypothesized that anti-CSF1R depletes both anti- and pro-inflammatory macrophage populations, whereas QPCTL selectively mediates the recruitment of monocytes that inhibit tumor immunity.

**Fig. 4.**
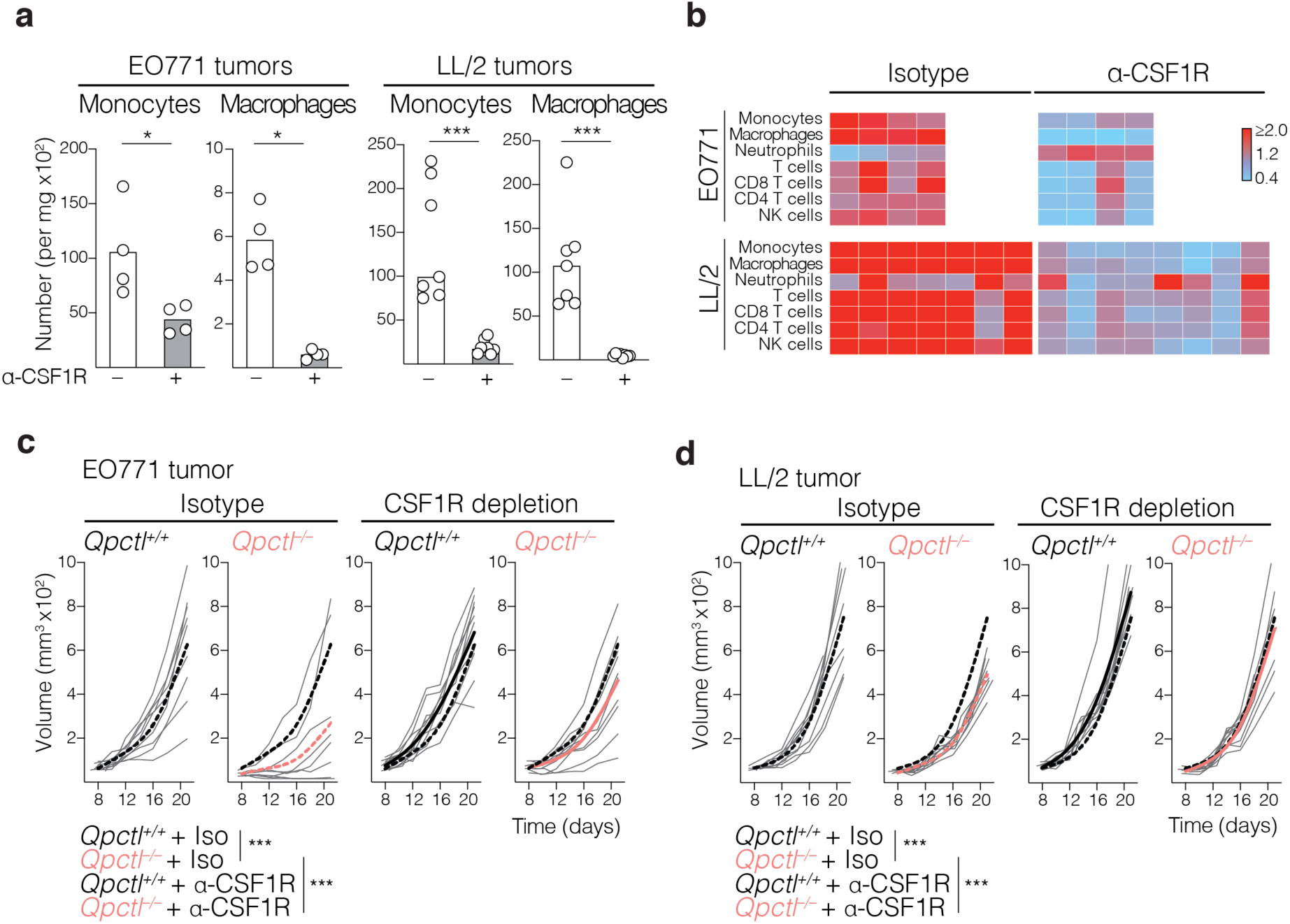
Anti-CSF1R treatment reverses the beneficial effect of *Qpctl*-loss on tumor growth. **a,b,** WT mice were inoculated with EO771 or LL/2 tumors and treated with ctrl isotype or anti-CSF1R antibody. **a**, Quantification of infiltrating leukocytes in tumors excited at day 14 (EO771) or 19 (LL/2) after tumor inoculation is shown. **b**, Heat map represents fold change leukocyte number, relative to median values of ctrl groups (n = 8 (EO771) or 15 (LL/2) mice). Each column represents a mouse. **c,** *Qpctl^−/−^* and *Qpctl^+/+^* littermate mice were inoculated with *Qpctl^−/−^* or *Qpctl^+/+^* EO771 cells, respectively, and treated with ctrl isotype or anti-CSF1R antibody. Tumor growth was measured over time. Gray lines represent individual mice. Overlay of fitting spline curves for each group is shown. Black dotted lines represent spline curves from ctrl *Qpctl^+/+^* groups (n = 9 (isotype) or 10 (anti-CSF1R) mice). **d,** *Qpctl^−/−^* and *Qpctl^+/+^* littermate mice were inoculated with LL/2 tumors and treated as described in (**c**). Tumor growth was measured over time (n = 10 mice per group). Bars are medians and symbols individual mice. Data shown is representative of one experiment. All experiments were repeated independently (⩾2 times). * *p*⩽0.05, *** *p*⩽0.001. P values are from nonparametric Mann-Whitney test (**a**) or Two-way ANOVA test (**c**,**d**).

### Modulation of QPCTL reshapes the composition of the intratumoral myeloid infiltrate

To characterize the remodeling of the monocyte/macrophage compartments in the absence of *Qpctl* expression, we performed single cell RNA-Seq analysis on CD45^+^ cells (leukocytes) infiltrating WT and *Qpctl^−/−^* LL/2 tumors, recovering an average of 2500 cells per sample (**Extended Data Fig. 8a**). Unsupervised clustering using Uniform Manifold Approximation and Projection (UMAP) identified five clusters: Mono/Mac/DCs (monocytes, macrophages and dendritic cells) characterized by expression of *Lyz2, Apoe* and MHCII components; Neutrophils, expressing *S100a8/9* and *Cxcl2*; Lymphocytes, identified by expression of *Gzmb* and *Tcrbc;* B cells and other non-hematopoietic cells (**Extended Data Fig. 8b–d**).

After removing neutrophils and non-myeloid cells, and re-clustering, the Mono/Mac/DCs could be assigned to 10 distinct clusters (**Fig. 5a**). We identified three clusters of monocytes: Mono_Ly6C, Mono_Ccr1/2 and Mono_Cx3cr1, all expressing *Csf1r*, *Ccr2* and *Ly6c2* (**Fig. 5b and Extended Data Fig. 9a-c**). All three monocyte populations expressed a gene signature similar to circulating monocytes ^34^ (**Extended Data Fig. 9d and Supplementary Table 1**). In addition, we identified five macrophage clusters characterized by expression of *Csf1r* and *Apoe*, and low *Ly6c2*: Mac_Ccr2; Mac_Cxcl2; Mac_MHCII; Mac_Timp2 and Mac_Ki67 (**Fig. 5b, Extended Data Fig. 9a,c**). We also identified two clusters of classic dendritic cells (cDCs) with high expression of MHC class II molecules and *Dpp4*, segregated by their expression of *Ccr7* and *Ccl22* (cDC1) or *Cd209a* and *Klrd1* (cDC2) (**Fig. 5b and Extended Data Fig. 9a,e**).

**Fig. 5.**
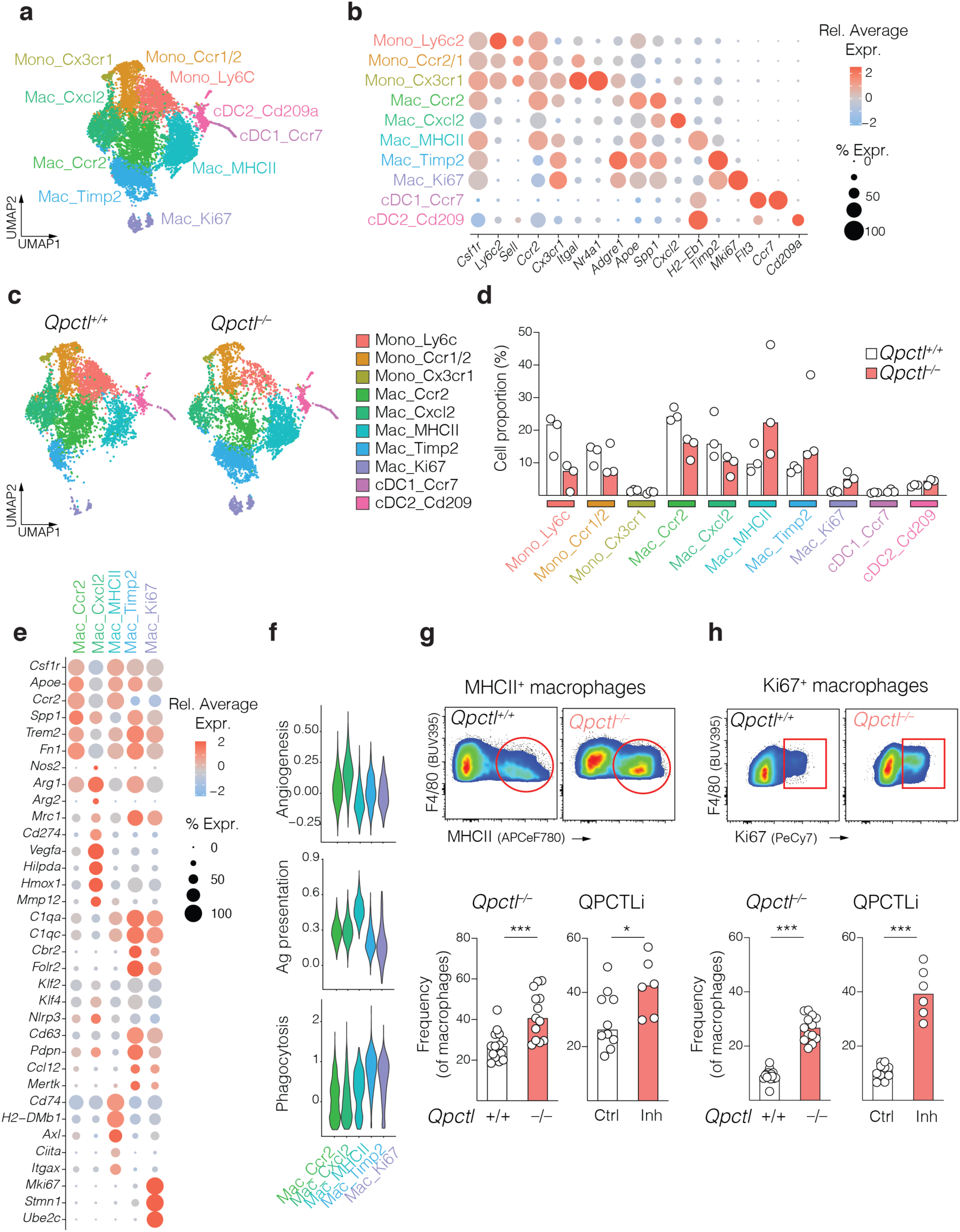
Modulation of QPCTL reshapes the composition of the intratumoral myeloid cell infiltrate. **a**,**b**,**c**,**d**,**e**,**f**, *Qpctl^−/−^* and *Qpctl^+/+^* littermate mice were inoculated with *Qpctl^−/−^* or *Qpctl^+/+^* LL/2 tumors, respectively. On day 14 after tumor inoculation, tumors were excised and processed for single cell RNAseq. **a**, UMAP plots of Mo/Mac/DCs clusters, from merged samples (n = 6, 3 samples per genotype). **b**, Dot plots of selected markers from merged samples. Dot size indicates the proportion of cells in each cluster expressing a gene and color shading indicates the relative level of gene expression. **c**, UMAP plots of Mo/Mac/DCs clusters, segregated by genotype (n = 3 samples per genotype). **d**, Abundance of clusters within *Qpctl^−/−^* and *Qpctl^+/+^* cells. **e**, Dot plots of selected markers from merged samples. **f**, Violin plots representing the expression of indicated gene signatures across macrophage clusters. **g**,**h**, LL/2 tumors were analyzed by flow cytometry. **g**, Representative flow cytometry plots identifying MHCII^+^ macrophages (red circle) among tumor infiltrating macrophages. Bar graph representing quantification of MHCII^+^ macrophages is shown (n= 15 or 13 (*Qpctl^−/−^*) mice). Right graph represents quantification of MHCII^+^ macrophages infiltrating LL/2 tumors inoculated in WT mice, treated with ctrl or QPCTLi (n = 10 or 6 (QPCTLi) mice). **h**, Identification and quantification of Ki67^+^ macrophages in samples described in (**g**). Bars are medians and symbols individual mice. Data shown is representative of one experiment or pooled from 2 experiments (**g**,**h**, left graphs). Single cell RNAseq experiment was done once, with 3 biological replicates per genotype. \**p*⩽0.05, \*\*\**p*⩽0.001. P values are from nonparametric Mann-Whitney test.

We next assessed the relative contribution of either genotype to each cluster, and found that *Qpctl^−/−^* cells were less abundant within the monocyte clusters (**Fig. 5c,d**), confirming our flow cytometric analysis. Additionally, *Qpctl^−/−^* cells were underrepresented in two macrophage subsets, Mac_CCR2 (*Qpctl^−/−^* cells 37%) and Mac_Cxcl2 (*Qpctl^−/−^* cells 35%) (**Fig. 5c,d**). Mac_CCR2 macrophages expressed markers of immunosuppression and alternative macrophage activation, such as Osteopontin, *Spp1* and Fibronectin, *Fn1* ^17^, whereas Mac_Cxcl2 macrophages presented an angiogenic signature, with high expression of the angiogenic factor, *Vegfa,* matrix metallopeptidase 12, *Mmp12* and hypoxia-related genes such as *Hilpda* and *Hmox1* ^35^ (**Fig. 5e,f and Extended Data Fig. 9a,f and Supplementary Table 1**). Mac_Cxcl2 also expressed the highest levels of the classic immunosuppressive genes *Cd274* (PD-L1) *Arg1* and *Arg2* (**Fig. 5e and Extended Data Fig. 9f**).

In contrast, loss of *Qpctl* resulted in a relative enrichment of the macrophage cluster Mac_MHCII (70% *Qpctl^−/−^* cells**, Fig. 5c,d**), characterized by a high expression of antigen presentation components (*H2-Eb1*, *H2-Ab1*, *H2-A*, *H2-DMb1*, *H2-DMa*, *Ciita* and *Cd74,* **(Fig. 5e,f, Extended Data Fig. 9a,f and Supplementary Table 1**), suggesting a role in T cell activation. We confirmed the increase in MHCII- expressing macrophages in *Qpctl^−/−^* and QPCTLi-treated LL/2 tumors using flow cytometry **(Fig. 5g**). *Qpctl^−/−^* cells were also enriched in Mac_Timp2 (**Fig. 5c,d**), characterized by expression of the tissue inhibitor of metalloproteinases 2,*Timp2* (**Fig. 5b**). The vast majority of proliferating macrophages Mac_Ki67, expressing cell cycle genes (*Mki67, Stmn1, Ube2c*), were also of *Qpctl^−/−^* genotype (**Fig. 5c-e**). Using flow cytometry, we confirmed that deletion or inhibition of QPCTL induced proliferation of intra-tumoral macrophages (**Fig. 5h**). SingleR analysis demonstrated that Mac_Ki67 exhibited a high degree of similarity with Mac_Timp2 (**Extended Data Fig. 9g**), suggesting that the Mac_Ki67 were mainly proliferating Mac_Timp2 macrophages. Mac_Timp2 and Mac_Ki67 macrophages expressed a phagocytosis gene signature (**Fig. 5f and Supplementary Table 1**) as well as *Klf2*, *Klf4* (previously identified as important for phagocytosis of apoptotic cells and proliferative properties of tissue-resident macrophages ^30, 36–38^) and *Cbr2* and *Folr2*, which are overexpressed in tissue-resident macrophages ^39^ (**Fig. 5e**). The lack of *Ccr2* expression further supported the notion that the F4/80^hi^ (*Adgre1*) Mac_Timp2 and Mac_Ki67 macrophages did not differentiated from recently infiltrating monocytes ^40^ (**Fig. 5b and Extended Data Fig. 9f**). These data do not exclude that Mac_Timp2 and Mac_Ki67 macrophages might have originated from monocytes, which have been proposed to replace embryonically established macrophages niches during homeostasis, adapting a phenotype distinct from circulating monocytes^40^. Thus, loss of *Qpctl* resulted in the reduction and a concomitant expansion of distinct populations of tumor-associated macrophages explaining the overall similar number of macrophages when comparing WT and *Qpctl^−/−^* tumors by flow cytometry. Of note, we did not find a clear association between the canonical M1 and M2 signatures with any of the macrophage populations^41^ (**Extended Data Fig. 9h and Supplementary Table 1**).

The relative expansion of F4/80^hi^ and Ki67^+^ macrophages following *Qpctl* loss contrasted with the preferential elimination of these populations upon anti-CSF1R treatment, as recently described ^17^. We scored our macrophage populations against gene signatures defined by Zhang and colleagues, and found that monocyte-derived Mac_Cxcl2 macrophages, which are diminished in the absence of *Qpctl* expression, were similar to the described Macro_Vegfa population (**Extended Data Fig. 9i**), characterized by the expression of pro-angiogenic genes and spared by anti-CSF1R treatment ^17^. Interestingly, this population expressed the lowest levels of *Csf1r* (**Extended Data Fig. 9j**), partially explaining their resistance to anti-CSF1R treatment.

Taken together, our data suggested that loss of *Qpctl* induced remodeling of the monocyte/macrophage compartments in the tumor microenvironment, characterized by an enrichment of tumor-associated macrophages with potential phagocytic properties, and macrophages with high expression of genes involved in antigen-presentation. In addition, loss of *Qpctl* impaired the accumulation of macrophages with putative immunosuppressive and pro-angiogenic functions contrasting, once more, to anti-CSF1R treatment.

### Loss of *Qpctl* in the MCP-producing cells is sufficient for inhibition of tumor growth

While it is challenging to genetically manipulate specific macrophage subsets, we investigated the mechanism(s) by which QPCTL regulates anti-tumor responses. Besides CCL2 and CCL7, QPCTL has many substrates which could impact tumor growth ^42, 43^. Notably, it was recently reported that loss of QPCTL in tumor cells results in diminished N-terminal modification of the “don’t-eat-me” signal CD47, thus abrogating engagement of SIPRα and enhancing phagocytic uptake by macrophages ^44, 45^. To address if loss of *Qpctl* in tumor cells was the main contributing factor in our models, we inoculated WT and *Qpctl^−/−^* mice with either WT or *Qpctl^−/−^* tumors. We found that loss of host-derived *Qpctl* was sufficient to inhibit CCL7 cyclization and attenuate tumor growth in the EO771 model **(Fig. 6a–c)**, thus ruling out a critical contribution of the QPCTL/CD47 axis in this model.

**Fig. 6.**
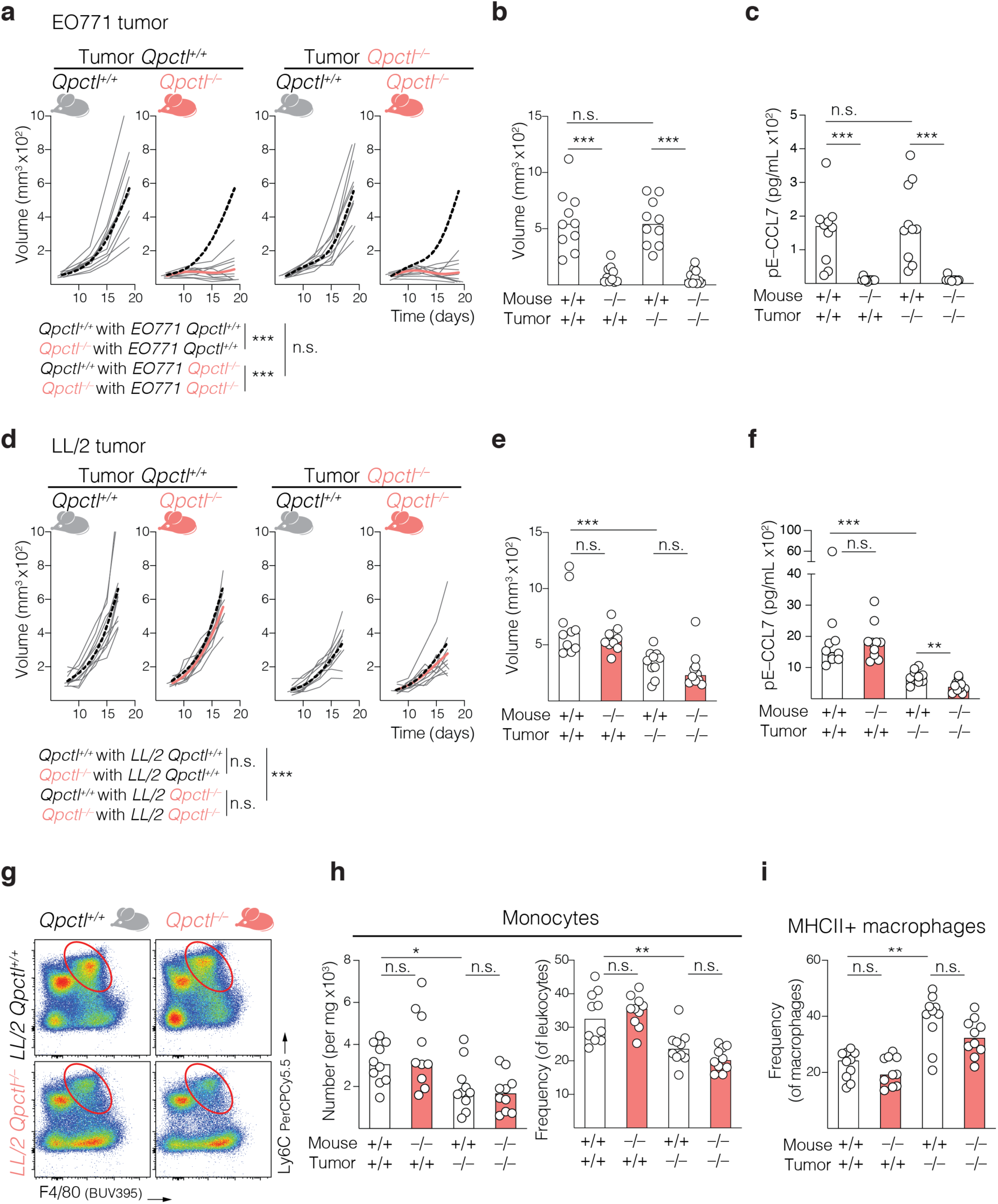
Loss of *Qpctl* in the MCP-producing cells is sufficient for inhibition of tumor growth. **a**,**b**,**c**, *Qpctl^−/−^* and *Qpctl^+/+^* littermate mice were inoculated *Qpctl^−/−^* or *Qpctl^+/+^* EO771 tumors, respectively. **a**, Tumor growth was measured over time (n = 10 mice per group). Gray lines represent individual mice and overlay of fitting spline cubic curves for each group is shown. Black dotted lines represent spline curves from *Qpctl^+/+^* host groups. **b**, Bar graph depicting tumor volume at the end of the study is shown. **c**, Quantification of CCL7 cyclisation in plasma from mice inoculated with EO771 tumors. **d**,**e**,**f**,**g**,**h**,**i**, *Qpctl^−/−^* and *Qpctl^+/+^* littermate mice were inoculated with *Qpctl^−/−^* or *Qpctl^+/+^* LL/2 tumors, respectively. **d**, Tumor growth was measured over time. Gray lines represent individual mice and overlay of fitting spline cubic curves for each group is shown (n = 10 mice per group). Black dotted lines represent spline curves from control *Qpctl^+/+^* host groups. **e**, Bar graph depicting tumor volume at the end of the study is shown. **f**, Quantification of CCL7 cyclisation in plasma from mice inoculated with LL/2 tumors. **g**, Tumors were excised 14 days after inoculation and analyzed by flow cytometry. Representative flow cytometry plots identifying Ly6C^+^ monocytic cells (red circle) among tumor infiltrating leukocytes. **h**, Number and frequency of tumor infiltrating Ly6C^+^ monocytic cells are depicted (n = 10 mice per group). **i**, Quantification of MHC class II^+^ macrophages is shown (n = 10 mice per group). Bars are medians and symbols individual mice. Data shown is representative of one experiment (**a**-**f**) or pooled from 2 experiments (**h**,**i**). All experiments were repeated independently (⩾3 times). ns, not significant, \**p*⩽0.05, \*\**p*⩽0.01, *** *p*⩽0.001. P values are from nonparametric Mann-Whitney test (**b**,**c**,**e**,**f**,**h**,**i**) or Two-way ANOVA test (**a**,**d**).

By contrast, WT LL/2 cells grew similarly in WT or *Qpctl^−/−^ mice*, and faster than *Qpctl^−/−^* LL/2 cells, implanted in either mouse strains **(Fig. 6d–e)**, demonstrating that loss of *Qpctl* in LL/2 tumor cells was the determinant to limit their growth *in vivo*. While CD47 may in part contribute to tumor clearance in this model, we found that LL/2 cells were the main producers of MCPs and determined the cyclization of CCL7 (**Fig. 6f**). Supporting our interpretation, WT mice challenged with *Ccl2^−/−^Ccl7^−/−^* LL/2 tumors (generated using CRISPR/Cas9, **Extended Data Fig. 10a**) had reduced levels of plasma chemokines (**Extended Data Fig. 10b)**, and showed reduced tumor growth (**Extended Data Fig. 10c,d)**. Thus, *Qpctl* expression acts in the cells producing the MCPs, and loss of either QPCTL or MCPs is associated with tumor growth inhibition.

We further demonstrated that loss of *Qpctl* in LL/2 cells was sufficient to limit monocyte infiltration in tumors (**Fig. 6g,h**), and to increase the frequency of intratumoral MHCII^+^ macrophages (**Fig. 6i**). Additionally, loss of *Ccl2* and *Ccl7* in the tumor cells also reduced infiltration of Ly6C^+^ monocytic cells, without impacting the number of tumor-associated macrophages (**Extended Data Fig. 10e,f**), but increasing the proportion of MHCII^+^ macrophages, similarly to the result obtained upon deletion of *Qpctl* (**Extended Data Fig. 10g**).

Together, these findings demonstrated that inhibition of MCP function, and the resulting reduction in monocyte infiltration, correlates with tumor growth inhibition following *Qpctl*-deletion in our models. Thus, targeting QPCTL engages multiple myeloid anti-tumor mechanisms, in addition to the described function in facilitating tumor cell phagocytosis^44, 45^.

### Loss or inhibition of QPCTL synergizes with anti-PD-L1 to increase CD8^+^ T cell responses

Our data showed that loss of *Qpctl* reshaped the macrophage compartment in tumors in favor of populations with a pro-inflammatory profile. The comparison with anti-CSF1R treatment had further shown that these macrophage populations were required to sustain a lymphocyte infiltrate. We therefore assessed if CD8^+^ T lymphocytes contributed to the tumor growth inhibition observed with *Qpctl* loss. We selected the EO771 model, which is characterized by a larger T cell infiltrate compared to LL/2 tumors (**Extended Data Fig. 6b**), inoculated WT or *Qpctl^−/−^* mice with WT or *Qpctl^−/−^* tumors, respectively, and depleted CD8^+^ T lymphocytes using an anti-CD8a antibody (**Extended Data Fig.10h**). CD8^+^ T cell depletion in WT mice accelerated tumor growth, demonstrating that CD8^+^ T cells played a role in the anti-tumor response in this model (**Fig. 7a**). CD8^+^ T cell depletion in *Qpctl^−/−^* mice restored tumor growth (**Fig. 7a**), establishing that tumor growth inhibition following *Qpctl* loss required a CD8^+^ T cell response.

**Fig. 7.**
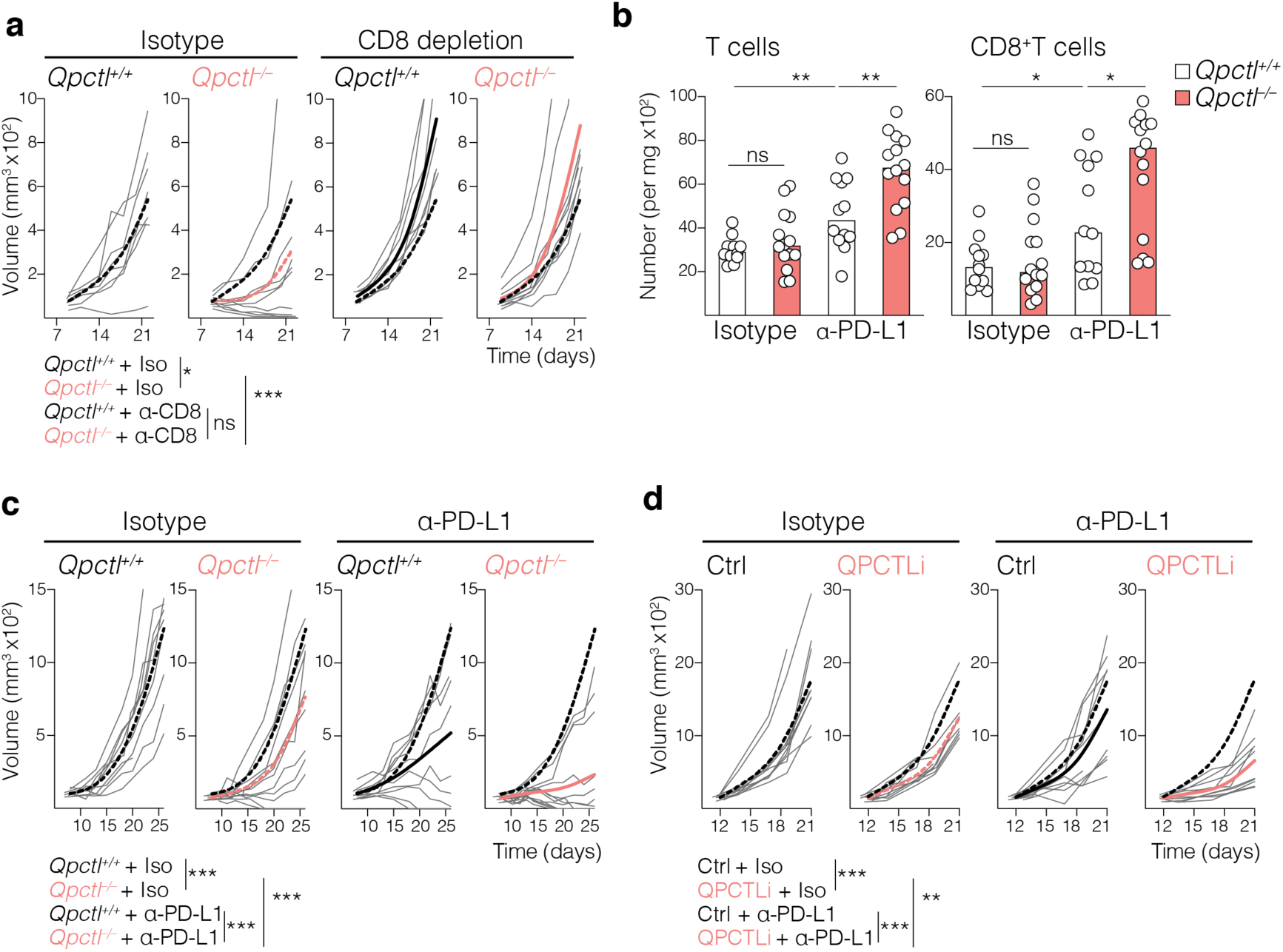
Loss or inhibition of QPCTL synergizes with anti-PD-L1 treatment to increase CD8^+^ T cell responses. **a**, *Qpctl^+/+^* and *Qpctl^−/−^* littermate mice were inoculated with *Qpctl^+/+^* or *Qpctl^−/−^* EO771 tumor cells, respectively. Mice were treated with ctrl isotype or anti-CD8 depletion antibody. Tumor growth was measured over time (n = 8 (WT) or 10 (*Qpctl^−/−^*) mice per group). Gray lines represent individual mice. Overlay of fitting spline curves for each group is shown. Black dotted lines represent spline curves from the isotype treated *Qpctl^+/+^* group. **b**, *Qpctl^+/+^* and *Qpctl^−/−^* littermate mice were inoculated with *Qpctl^+/+^* or *Qpctl^−/−^* EO771 tumor cells, respectively. On day 9 after tumor cell inoculation, mice received ctrl isotype or anti-PD-L1 blocking antibody. Tumors were excited 7 days after initiation of treatment and analyzed by flow cytometry. Number of tumor-infiltrating T cells and CD8^+^ T cells is shown (n = 12 (WT) or 14 (*Qpctl^−/−^*) mice per group). **c**, *Qpctl^+/+^* and *Qpctl^−/−^* littermate mice were inoculated as described in (**b**). On day 12 after tumor cell inoculation, mice were treated with control isotype or anti-PD-L1 blocking antibody. Tumor growth was measured over time. Gray lines represent individual mice. Overlay of fitting spline curves for each group is shown. Black dotted lines represent spline curves from the isotype treated *Qpctl^+/+^* group (n = 10 (WT) or 12 (*Qpctl^−/−^*) mice per group). **d**, WT mice were inoculated with EO771 cells. On day 13 after tumor cell inoculation, mice were treated with ctrl isotype or anti-PD-L1 blocking antibody. In parallel, mice started treatment with QPCTLi or ctrl vehicle. Tumor growth was measured over time. Gray lines represent individual mice. Overlay of fitting spline curves for each group is shown. Black dotted lines represent spline curves from the ctrl vehicle and isotype treated group (n = 10 mice per group). Bars are medians and symbols individual mice. Data shown is representative of one experiment (**a**,**d**), or pooled from 2 (**b**,**c**). All experiments were repeated independently (⩾2 times). * *p*⩽0.05; ** *p*⩽0.01 *** *p*⩽0.001. P values are from nonparametric Mann-Whitney test (**b**), Two-way ANOVA test (**a**,**c**,**d**).

Because we had not observed differential accumulation of T cells upon *Qpctl* deletion (**Extended Data Fig. 7a**), we next assessed the potential for a combination effect with a T cell-targeted treatment, such as checkpoint inhibition. We inoculated WT or *Qpctl^−/−^* mice with WT or *Qpctl^−/−^* EO771 tumors, and treated them with an isotype control or anti-PD-L1 antibody. Treatment with anti-PD-L1 increased the number of tumor-infiltrating CD8^+^ T cells in WT mice and even more in the *Qpctl* -deficient setting, whereas *Qpctl* loss by itself did not affect CD8^+^ T cell numbers (**Fig. 7b**). Granzyme B production by CD8^+^ T cells was not altered across the four groups (**Extended Data Fig. 10i**). WT tumors treated with anti-PD-L1 grew slower than isotype-treated tumors, and further tumor growth inhibition was observed with anti-PD-L1 in *Qpctl*-deficient tumors (**Fig. 7c**). To test if therapeutic modulation of QPCTL also synergized with anti-PD-L1, we treated mice with established EO771 tumors with QPCTLi, anti-PD-L1 or the combination of both. In this therapeutic setting, we observed that the combination of anti-PD-L1 with QPCTLi resulted in further attenuation of tumor growth, when compared to control isotype and single treatment groups (**Fig. 7d**).

Taken together, our data demonstrated that genetic loss or inhibition of QPCTL synergizes with checkpoint blockade to enhance T cell functions and to limit tumor growth.

## DISCUSSION

Targeting of chemokines has long been pursued as a potential strategy for modulating cellular trafficking in different disease settings ^46^. Previous work revealed that modulation of enzymes that modify the N-terminus of chemokines, such as the extracellular peptidase DPP4, constitute effective targets to potentiate immune cell migration and anti-tumor immunity ^22, 23^. Here, we demonstrated that the main monocyte chemoattractants CCL2 and CCL7 are insensitive to DPP4-inactivation *in vivo* because of an intracellular mechanism of N-terminal cyclization mediated by the Golgi-associated enzyme QPCTL. This pathway is essential to ensure the homeostatic functions of MCPs as mediators of monocyte export from the bone marrow, and represents a mode of protection that distinguishes MCPs from other inflammatory chemokines, such as CXCL10.

In addition to its role in homeostatic monocyte migration, our work also revealed that QPCTL is a critical regulator of monocyte migration into solid tumors. When infiltrating tumors, monocytes originate diverse myeloid cell populations, adapting to niche-specific signs and inflammation patterns ^47^. High-resolution analyses revealed that healthy-tissue macrophages may also be precursors of tumor myeloid cells, further adding to the complexity of the myeloid compartment observed across tumors ^11^. Our study demonstrated that *Qpctl* loss induces profound reshaping of the myeloid cell composition in tumors, with a decrease in *Ccr2*-expressing cells, including *Ly6c2*-expressing monocytes and two subsets of macrophages. Curiously, a third population of *Ccr2*-expressing macrophages (Mac_MHCII) was rather enhanced upon loss of *Qpctl*. This population expressed genes related to antigen presentation, suggesting a pro-inflammatory role in T cell interaction and activation. Previous studies suggested that MCP signaling influences macrophage polarization ^48, 49^. Notably, our studies using *Ccl2^−/−^Ccl7^−/−^* tumors also revealed modulation of macrophage polarization, with increased MHC II expression. It is likely that, besides impairing chemoattraction, modulation of QPCTL also influences other functions triggered by chemokine signaling (or other putative QPCTL targets), polarizing the remaining *Ccr2*-expressing cells in the tumor.

Loss of QPCTL resulted in a striking accumulation of two macrophage populations expressing high levels of F4/80 and lacking CCR2 expression. Besides markers of tissue macrophages, these subsets expressed gene signatures of phagocytic activity and cell cycling, indicating proliferative potential ^30, 37^. The role of monocyte-derived *versus* tissue macrophage-derived myeloid cells in tumor immunity has been the center of active discussion but remains largely elusive. The discrepancies in functions attributed to myeloid cells of different ontogenies across different studies and models suggest that the tissue microenvironment may influence their activity in tumors ^3, 50^. Ultimately, a comprehensive understanding of tumor macrophage functions depends on our ability to selectively target and study individual populations. Our study demonstrates that QPCTL targeting constitutes an effective approach to preferentially impair monocyte entry into the tumors, creating a unique opportunity to study the role of macrophages of different origins.

The important anti-tumor properties of macrophages are highlighted by the poor pre-clinical and clinical activity of approaches that broadly eliminate myeloid cells, such as anti-CSF1R agents ^13, 14^. Our results further emphasized the putative role of macrophage populations in controlling tumor growth. Recent advances in understanding the effects of anti-CSF1R at the single cell level revealed that this treatment eliminates certain macrophage populations preferentially while sparing others, a result previously overlooked when using conventional immunophenotyping methods. When comparing deletion of *Qpctl* with anti-CSF1R treatment, we found that a macrophage population characterized by high expression of pro-angiogenic and hypoxic genes remained after anti-CSF1R treatment, but was diminished in *Qpctl^−/−^* conditions. In sharp contrast, anti-CSF1R treatment preferentially eliminated F4/80^hi^ macrophages with proliferative capacity, which were expanded upon loss of *Qpctl*. As CSF1R targeting did not result in any benefit in our models, it is tempting to speculate that F4/80^hi^ tissue macrophages have anti-tumor properties revealed by *Qpctl* loss. It remains to be defined, however, how incoming monocytes influence tissue macrophage biology in tumors. Additionally, further studies are needed to understand the specific role(s) of macrophages in anti-tumor responses elicited by loss of *Qpctl*. High resolution analysis of myeloid cell modulators in tumors, paired with tumor growth data remains a powerful way to further understand the interplay between monocyte/macrophage populations and their distinct functions.

In addition to CCL2 and CCL7, both QPCT and QPCTL have other substrates, some of which might impact inflammation and sensitivity to immune cell recognition ^42, 43^. Notably, N-terminal cyclization is important for CD47 function, which desensitizes macrophage-mediated phagocytosis of tumor cells ^44, 45^. *Qpctl* knockout induced accumulation of macrophages expressing phagocytic genes, suggesting that this modality of anti-tumor immunity may be enhanced via two QPCTL-mediated mechanisms and highlighting the potential of targeting QPCTL as a means of engaging and enhancing multiple anti-tumor mechanisms, such as macrophage-mediated phagocytosis and adaptive T cell responses. Besides CD47, the human monocyte chemoattractants CCL13, CCL15, CCL16 and CX3CL1 also possess a glutamine as N-terminal residue and may be modified (and protected) by QPCTL ^43^. Targeting QPCTL represents, therefore, a robust means to prevent monocyte migration across inflammatory conditions, as it overcomes the high redundancy existing in the chemokine family, which is a limitation for strategies targeting individual chemokines or chemokine/receptors.

Glutaminyl cyclase inhibitors are being developed for the treatment of Alzheimer’s disease, to prevent the generation of pathogenic pyroglutamate-modified proteins ^32, 33^. Our study validates that QPCTLi recapitulates many of the phenotypes observed using models of genetic *Qpctl* loss. Pharmacological QPCTLi prevents monocyte accumulation in established tumors, promoting remodeling of the macrophage compartments and enhancing anti-tumor activity of anti-PD-L1. Our findings suggest that QPCTLi could be used in combination with various immunomodulatory agents, such as checkpoint inhibitors, or tumor-targeted agents to enhance antibody-mediated phagocytosis of tumor cells.

## METHODS

### Mice and ethics statement

Wild-type (WT) C57BL/6J or C57BL/6N CD45.2 and C57BL/6 CD45.1 mice were obtained from Jackson Laboratory or Charles Rivers, USA. WT CD45.1/CD45.2 and *Dpp4^−/−^* mice were bred in Genentech. *Qpct^− /−^* and *Qpctl^−/−^* mice were generated in Genentech, following homologous recombination and mouse embryonic stem cell (ES) technology. For the *Qpct*^−/−^ strain, a gene targeting vector was constructed with a 2040 bp 5’ arm of homology corresponding to GRCm38/mm10 chr17: 79,050,226 – 79,052,265 and a 2151 bp arm of 3’ homology arm corresponding to chr17:79,055,051-79,057,201. Exon1 was deleted after the ATG plus 20bp of intron corresponding to chr17:79,052,269-79,052,408. For the *Qpctl^−/−^* strain, a gene targeting vector was constructed with a 2060 bp 5’ arm of homology corresponds to GRCm38/mm10 chr7: 19,149,183-19,151,242 and a 1976 bp arm of 3’ homology arm corresponds to chr7:19,146,990-19,148,965. Exon1 was deleted after the ATG plus 10bp of intron corresponding to chr7:19,148,966-19,149,182. Egfp-SV40-FRT-pgk-neo-FRT right after the ATG of exon1 in both strains. The final vector was confirmed by DNA sequencing, linearized and used to target C2 (C57BL/6N) ES cells using standard methods (G418 positive and ganciclovir negative selection) C57BL/6N C2 ES cells were electroporated with 20 ug of linearized targeting vector DNA and cultured under drug selection. Positive clones were identified using long range PCR followed by sequence confirmation. Correctly targeted ES cells were subjected to karyotyping. Euploid gene-targeted ES cell clones were treated with Adeno-FLP to remove PGK neomycin and ES cell clones were tested to identify clones with no copies of the PGK neomycin cassette and the correct sequence of the targeted allele was verified. The presence of the Y chromosome was verified before microinjection into albino BALB/c embryos. Germline transmission was obtained after crossing resulting chimeras with C57BL/6N females. Genomic DNA from pups born was screened by long range PCR to verify the desired gene targeted structure before mouse colony expansion. For genotyping the *Qpct^−/−^* strain, the following primers were used: 5′-GGCAGACACAATCAATCC-3’, 5’-CAGCAGGTGGAGAGTG-3’ and 5′-TTACGTCGCCGTCCA-3’ amplified 164-bp wild-type and 186-bp knock-in DNA fragments.7-12 weeks old female mice were used. For genotyping the *Qpctl^−/−^* strain the following primers were used: 5′-ACAGCCAATCGCAATCG -3’, 5’-CACGCCGTAGGTCAGG -3’ and 5′-GATATAGAAAGCCAAGCCCATA-3’ amplified 287-bp wild-type and 341-bp knock-in DNA fragments. *Qpct^−/−^Qpctl^−/−^ and Dpp4^−/−^Qpctl^−/−^ double* mutant mice were generated by crossing single mutant strains. To generate chimeric mice, bone marrow cells from WT mice (CD45.1) or *Qpctl^−/−^* mice (CD45.2) were collected from femurs of mice and transferred onto irradiated WT or *Qpctl^− /−^* mice. Irradiation of recipient mice was done following a protocol of 2 times 5.5Gy dosage, 6h apart from each other. Two million bone marrow cells were transferred via intravenous route to each recipient mouse, 24h after the first dose of irradiation. Bone marrow chimeras generated were 4 groups: WT (CD45.1) into WT (CD45.2); *Qpctl^−/−^* (CD45.2) into *Qpctl^−/−^* (CD45.2); WT (CD45.1) into *Qpctl^−/−^* (CD45.2) and *Qpctl^−/−^* (CD45.2) into WT (CD45.1 mice). In all experiments control WT (+/+) mice refer to mice bred within the same colony, and that do not carry the mutation (littermates). All mice were housed at Genentech in standard rodent micro-isolator cages and were acclimated to study conditions for at least 7 days before manipulation. Animals were 7-20 weeks old. All animal experimental protocols were approved by the Genentech institutional animal care and use committee, responsible for ethical compliance.

### Cell lines

The murine colon adenocarcinoma Lewis lung carcinoma LL/2 and the human monocytic cell line THP-1 were obtained from ATCC. The murine breast adenocarcinoma EO77 was from CHBiosystems. All cell lines were tested negative for *Mycoplasma*. Cell lines were kept at low passage and maintained in complete R10 media (RPMI1640 medium (Genentech) supplemented with 10% heat inactivated FBS (Sigma or VWR), 200mM L-Glutamine (Sigma); 100 units/mL of penicillin and 100 µg/mL of streptomycin (Gibco)). Generation of *Qpctl^−/−^* tumor lines was done via CRISPR / Cas9 technology using a crRNA:tracrRNA duplex guide RNA with specificity for the second exon of murine QPCTL. The tracrRNA (TTATTGATTGTGCGACCCCCGGG) and crRNA (IDT) were resuspended at a final concentration of 100µM in nuclease-free duplex buffer (IDT), at mixed at equimolar ratio, denatured for 5min at 95°C and allowed to cool down to room temperature. 3µl of assembled gRNA were mixed with 2µl (60pmol) of TruCut Cas9 protein V2 (Thermofisher) and incubated for 10min prior to nucleofection. 1.2 million cells were resuspendend in 20µl SE buffer (Lonza SE Cell Line 4D-NucleofectorTM X Kit S, #V4XC-1032), mixed with the gRNA/Cas9 cocktail and nucleofected in a Lonza 4D-Nucleofector using program CA137. Cells were then transferred into 180µl pre-warmed R10 media on a 96-well plate and let rest for 30min prior to transfer to a 6-well plate (2ml media per well). Cells were allowed to expand for several days before being tested. *Qpctl* knockout efficiency in bulk tumor cell lines was monitored by accessing the frequency of WT and mutant gene reads by sequencing (frequency of mutated gene reads was >95% for all lines, no sorting was performed) and assaying the capacity of the cells to produce and secrete N-terminal cyclized CCL7 (pE–CCL7) via SIMOA immunoassays. CRISPR / Cas9-mediated deletion of *Ccl2* and *Ccl7* in bulk tumor cell lines was done as described for *Qpctl* deletion. Deletion of chemokines was sequential using the following tracrRNAs: CACCAGTAGCCACTGTCCCCGGG and GCTGTTATAACTTCACCAATTGG for *Ccl7* and *Ccl2*, respectively. Efficacy of *Ccl7 and Ccl2* knockout was monitored by assaying the capacity of the cells to produce and secrete CCL7 and CCL2 via SIMOA immunoassays.

### Recombinant proteins

Recombinant mouse (m)CCL7 and human (h)CCL2 and hCCL7 were purchased from Peprotech. Recombinant hDPP4 was purchased from R&D. Recombinant mQPCTL was expressed in *E.coli* BL21(DE3) with an N-terminal His-tag, and purified by Ni-NTA followed by Superdex 75. His-tag was then cleaved by TEV protease and separated by Ni-NTA HiTrap SP HP. To generate pE’-CCL7, 12ug of mCCL7 were incubated with 10ug of mQPCTL, in a final volume of 50ul and incubated at 37 °C for 12-18h period. Truncated ΔQP’–hCCL7 was transiently expressed in HEK 293T cells using pRK5 vector with N-terminal His-FLAG tags, and purified by Ni Sepharose Excel (GE Healthcare) followed by Superdex 75 (GE Healthcare). Tags were removed by Enterokinase (Roche) treatment and purified away by Ni-NTA (Qiagen). Truncated ΔQP–hCCL2 was expressed in E. coli BL21(DE3) using the *E.coli* expression vector pET52b backbone, with N-terminal His-FLAG tags. Tags were removed by Enterokinase (Roche) treatment and purified away by Ni-NTA (Qiagen). ΔQP–mCCL7 was expressed in E. coli BL21(DE3) with N-terminal His-FLAG tags and purified by Ni-NTA followed by Superdex 75. Tags were removed by Enterokinase treatment and separated by HiTrap SP HP (GE Healthcare).

### SIMOA Immunoassays

Antibodies that recognize mCCL7 and mCCL2, independently of their N-terminal sequence were purchased from R&D (Cat# AF-456-NA and AF-479-NA, respectively). Antibodies recognizing specifically the cyclized isoforms of CCL7 and CCL2 (pE–CCL7 and pE–CCL2, respectively) or recognizing specifically truncated isoforms (ΔQP–CCL7 and ΔQP–CCL2) were produced at Genentech using a rabbit immune phage display technology. Rabbits were immunized with cyclized or truncated isoforms peptides. The variable regions of immunoglobulin heavy and light chains were amplified from splenocytes by PCR and used to make a single-chain Fv (scFv) phage display library on M13 phage. To obtain the cyclized isoform specific antibodies, the library was panned on cyclized peptides and counter-selected with the corresponding wild-type N-terminal peptides. Same panning strategy applied to the truncated isoforms specific antibody selection. Sequence unique phage clones that selectively bind to the cyclic or truncated isoforms were selected and subcloned into a mammalian IgG expression vector. Protein A-purified rabbit IgG were confirmed the selective binding activity by ELISA and SPR (surface plasmon resonance) with the corresponding peptides. Mouse CCL7 and CCL2 isoforms in serum, plasma and cell supernatants were monitored using SIMOA HD-1 assays (Quanterix). Homebrew SIMOA assays specific for total-chemokine or each chemokine modification (cyclized pE– and truncated ΔQP–) were produced in house per manufacturer’s instructions, using the antibodies previously referenced. Capture antibodies were AF457 (R&D, 0.2mg/mL) for total–CCL7; 1G2 (0.02mg/mL) for pE–CCL7; 2F3 (0.02mg/mL) for ΔQP– CCL7; AF479, (R&D, 0.1mg/mL) for total–CCL2; 1G2–VH02 (0.1mg/mL) for pE–CCL2 and 2G1 (0.2mg/mL) for ΔQP–CCL2. A biotinylated form of clone AF456 (R&D, 0.3 mg/mL) or clone MAB479 (R&D, 0.3 mg/mL) was used as detector antibody for all CCL7 and CCL2 assays, respectively. Plasma and serum samples were diluted in SIMOA Buffer (1xPBS, 2% BSA, 0.1%Tween, 5mM EDTA) at a 1:10 ratio for the total–CCL7 and pE–CCL7 assays, and 1:5 for the ΔQP–CCL7 assay. Dilutions were 1:2 for the total–CCL2 and pE–CCL2 assays, and 1:5 for the ΔQP–CCL2 assay. For evaluation of chemokine isoforms secreted from tumor cell lines *in vitro*, cells were grown to ∼80% confluence prior to media replacement with fresh media. Condition cell supernatants were analyzed 3 to 4h after media replacement.

### Flow cytometry

Antibodies for flow cytometry of mouse samples were: anti-CD3e (clone 145-2C11 in PECF594 from BD Biosciences Cat# 562286 or in FITC, from eBiosciences Cat# 11-0031-82, or clone 500A2 in AF700, from BD Biosciences Cat# 557984); anti-B220/CD45R (clone RA3-6B2 in APCeF780 from eBiosciences Cat# 47-0452-82, in AF700 from BD Biosciences Cat# 557957 or in FITC, from BioLegend Cat# 103206); anti-CD45 (clone 30-F11 in PECy7 from eBiosciences Cat# 25-0451-82 or in BV786, from BD Biosciences Cat# 564225); anti-CD45.1 (clone A20 in APC from BioLegend Cat# 110714); anti-CD45.2 (clone 104 in AF700 from eBiosciences Cat# 56-0454-82 or in APC from BioLegend Cat# 109814 or in pE-Cy7 from BioLegend Cat# 109830); anti-NK1.1 (clone PK136 in APCeF780 Cat# 47-5941-82 or APC cat# 17-5941-82 from eBiosciences, in AF700 from BD Biosciences Cat# 560515 or in FITC from BioLegend Cat# 108706); anti-CD11b (clone M1/70 in BV421, BD Biosciences Cat# 562605); anti-CD115/CSF1R (clone AFS98 in PECy7 from BioLegend Cat# 135524); anti-CCR2 (clone 475301 in APC from R&D systems Cat# FAB5538A); anti-CX3CR1 (clone SA011F11 in BV786 from Biolegend Cat# 149029); anti-F4/80 (clone T45-2342 in BUV395 from BD Biosciences Cat# 565614); anti-Ly6C (in PerCPCy5.5, clone HK1.4 from BioLegend Cat# 128012 or clone AL-21 from BD Biosciences Cat# 560525); anti-Ly6G (clone 1A8 in AF700, from BioLegend Cat# 127622); anti-MHC class II (I-A/I-E) (clone M5/114.15.2 in APCeF780 from eBiosciences Cat# 47-5321-82); anti-Siglec F (clone E50-2440 in PeCF594 from BD Biosciences Cat# 562757 or clone 1RNM44N in AF700 from Invitrogen Cat# 56-1702-82); anti-CD11c (clone N4.11 in PE, from eBiosciences Cat# 12-0114-82); Anti-Ki67 (clone SolA15 in PECy7 from Invitrogen Cat# 25-5698-82) and anti-Granzyme B (clone GB12 in APC, from Invitrogen Cat# MHGB05). For all flow cytometry stainings, cell suspensions were first incubated with mouse CD16/CD32 Fc–blocking antibodies (clone 2.4G2, BD Biosciences Cat# 553142) and Aqua LIVE/DEAD® Fixable Dead Cell Stain (Invitrogen, Carlsbad, CA) for 20 min at 4°C. Antibodies were diluted in PBS 1x and stainings were done for 20-30min at 4°C. Flow Cytometry data were collected with a BD LSRFortessa cell analyzer or FACSymphony (BD Biosciences, San Jose, CA) and analyzed using FlowJo Software (Version 10.2, FlowJo, LLC, Ashland, OR). For the determination of cell numbers, AccuCheck Counting Beads reagent (Invitrogen) was used.

### *In vivo* models of monocyte homeostasis and peritoneal inflammation

For evaluation of the role of DPP4 in monocyte homeostasis mice were fed with chow containing 1.1% of the DPP4 inhibitor (DPP4i) Sitagliptin (Envigo) ^22^ for up to 30 days. When indicated, mice were dosed (via intraperitoneal route) with 200ug of anti-CCL2 (clone 2H5, BioXcell) and 100ug of anti-CCL7 (polyclonal, R&D), or isotype controls (Armenian Hamster IgG and Goat IgG, respectively) ^51, 52^. Dosing started in parallel with administration of DPP4i, and was maintained for 30 days (biweekly dosing). For chemokine-mediated migration models, mice were intraperitoneally injected with sterile PBS or 1 μg of mCCL7 isoforms. Volume injected was 100ul per mouse. When indicated, mice were pre-treated with DPP4i. In adoptive cell transfer experiments, mice received an intravenous injection of 1×10^6^ monocytes before intraperitoneal injection with chemokine. Monocytes were isolated from bone marrow (femurs and tibias) with the EasySepTM Mouse Monocyte Isolation Kit (StemCell Technologies) according to the manufacturer’s protocol. Mice were sacrificed and peritoneal cells were collected in 8 ml of cold PBS, 16-18 h after injection. Cells were used for flow cytometry after spun down (250g, 5 min, 4°C). Only peritoneal lavages that were clear in color and did not present visible red blood cell contents (indicative of inaccurate intraperitoneal inoculation) were analyzed.

### *In vivo* tumor models

Mice were implanted subcutaneously with 0.1×10^6^ LL/2 cells resuspended in 100ul of Hanks’ Balanced Salt Solution (HBSS) and Matrigel (1:1 ratio). EO771 cells were implanted in female mice, in the 5^th^ mammary fat pad (0.1×10^6^, resuspended in 100ul of HBSS/Matrigel, 1:1 ratio). Anti-CSF1R (InvivoPlus, clone AFS98) or respective rat IgG2A isotype control (InvivoPlus, anti-trinitrophenol, 2A3) was dosed at 15 mg/kg IP starting on day -3 before tumor implantation followed by a twice a week regiment. For experiments using immune checkpoint blockers, mice with established tumors were randomized into groups and treated with either isotype (gp120) or anti-PD-L1 (clone 6E11) antibodies given at 10mg/Kg intravenously or the first dose, followed by 5mg/Kg intraperitoneally, thereafter twice a week. QPCTL inhibitor (QPCTLi, compound 11, WO2014140279) was administered at 50-75 mg/Kg orally, twice a day. For preventative studies, QPCTLi was administered to mice 1 to 4 days before tumor cell injection and for therapeutic studies, QPCTLi was administered to tumor bearing mice 7-13 days after tumor cell injection, as described in the figure legends. QPCTLi dosing was not interrupted during the study. MCT was administered to control groups. All groups receive an oral administration of 1-Aminobenzotriazole (ABT), an *in vivo* nonspecific inhibitor of cytochrome P450 enzymes at 50mg/Kg, once a day, 1 hour before administration of QPCTLi. When indicated, QPCTLi was administered to mice with established tumors, together with immune checkpoint blockade. For adoptive cell transfer experiments, LL/2 tumor bearing mice received an intravenous injection of 1.5×10^6^ WT CD45.1 congenic monocytes, 2 days before tumor analysis, which was performed at day 14 post-tumor cell inoculation. For CD8^+^ T cell depletion, rat anti- CD8 (ATCC-2.43) or control rat anti-GP120 (8G4) were administered intraperitoneally,10 mg/kg, starting at day -1 before inoculation of tumor cells, followed by two times per week. When indicated, mice were sacrificed and tumor-associate leukocyte infiltrates were evaluated by flow cytometry. Tumor growth was measured over time (2 to 3 times per week) with a caliper. Tumor volumes were measured in two dimensions (length and width) using Ultra Cal-IV calipers (Fred V. Fowler Co.; Newton, MA) and volume was calculated using the formula: Tumor size (mm3) = (length x width^2^) x 0.5.

### Tissue processing for cell suspensions

Blood samples (150ul) were collected in Eppendorf tubes containing 5ul of 0.25M EDTA, transferred to 5mL tubes containing 2mL of red blood cell (RBC) lysis buffer (ACK) and incubated for 5 min at room temperature (RT). After incubation, 2mL of PBS was added and samples were centrifuged (850g, 2min). Samples were subjected to another round of RBC lysis, with 1mL of ACK buffer. Bone marrow cells were extracted from one femur via centrifugation onto an Eppendorf tube (2500g, 2 min) resuspended in 1mL of ACK buffer for 5 min at RT, quenched with 2mL of PBS and spun down (850g 2min). Spleen and tumors were collected, cut in small pieces and digested in PBS supplemented with 0.2 mg/ml Collagenase P and 0.1mg/mL DNase I (both from Roche) for 20 min at 37°C. After incubation, digested tissues were transferred to 70-μm cell strainers placed onto 50mL falcon tubes. Digestion was quenched by adding PBS supplemented with 2% FCS and 5 mM EDTA (Gibco). Cell suspensions were obtained after filtration over the 70-μm cell strainer. Cell suspensions were resuspended in 2mL of ACK buffer for 5 min at RT, quenched with 5mL of PBS supplemented with 2% FCS and spun down at 250g 5min.

### Histology in mouse tissue

Tissues were fixed in 4% formalin for 24h and placed in 70% Ethanol for 24h, then paraffin embedded. Immunohistochemistry was performed on 4μm thick paraffin-embedded tissue sections mounted on Superfrost Plus glass slides. Unstained slides were baked at 70°C for 30 min. Sections were de-paraffinized and rehydrated to deionized water. For Iba-1 staining antigen retrieval was performed with 1X DAKO Target Retrieval Solution (pH 6) for 20 min at 99°C and cooled to 74°C. Subsequently, slides were rinsed with dH2O for 1-minute then rinsed with 1X Dako Wash Buffer (0.05M Tris, 0.15M NaCl, 0.05% Tween-20). Staining was performed on the Dako Autostainer (Agilent Technologies, Carpinteria, CA). Sections were incubated in mRabbit anti-Iba1, diluted to 0.3ug/mL in 3% Bovine Serum Albumin (BSA) in 1X PBS (GNE Media Prep, Cat. A4577) for 30-minutes. Detection was performed using PowerVision Poly-HRP Anti-Rabbit IgG Novocastra (Leica Biosystems, Cat. PV6119) for 30-minutes. Sections were counterstained, dehydrated in graded reagent alcohol [95% (1-minute x 2), 100% (1-minute x 2)] followed by clearing in xylene (1-minute x 3) and coverslipped with (xylene-based) permanent mounting medium (Sakura, 6419). For F4/80 staining, following deparaffinization and rehydration of sections, endogenous avidin and biotin were blocked with ScyTek Avidin Biotin Blocking Kit at room temperature per manufacturer’s directions then rinsed. Sections were blocked with 10% normal rabbit serum in 3% BSA/PBS for 30 minutes then incubated with anti-mouse F4/80 antibody diluted to 10ug/ml in blocking buffer for 60 minutes at room temperature. Sections were then incubated with biotinylated rabbit anti-rat antibody, diluted to 2.5 ug/ml in blocking buffer for 30 minutes, rinsed and incubated with diluted ABC Elite Reagent (Vector, PK-6100) for 30 minutes. Sections were then counterstained with Mayer’s hematoxylin, rinsed and cover slipped.

### Measurement of glutaminyl cyclase activity in mouse samples

For the *in vitro* reactions, 50 uL of mouse serum samples were mixed with 50 uL of 500 nM human CCL2 in 10 mM Tris pH 8 buffer and incubated at 37°C for varying times. CCL2 was affinity purified from the in vitro reactions using Phytip streptavidin coated tips (Phynexus) bound to biotinylated anti-CCL2 antibody (Invitrogen) using the Phytip protocol. 10 uL of affinity purified CCL2 is injected onto a Acquity BEH C18 column (Waters, 2.1mm x 50mm) and eluted over a gradient of 0% to 80% acetonitrile with 0.1% formic acid. Q and pE forms of CCL2 were quantitated using Multiple Reaction Monitoring (MRM) with a QTRAP 6500 triple quad mass spectrometer (Sciex). For the Q–CCL2, Q1 mass = 965.4, Q3 mass = 1204, collision energy = 35V, declustering potential = 45. For the pE–CCL2, Q1 mass = 1086, Q3 mass = 660.7, collision energy = 45V, declustering potential = 45. Source parameters used were curtain gas = 25, collision gas = medium, IonSpray voltage = 5500, temperature = 650, ion source gas 1 and 2 = 60. Percent conversion of Q–CCL2 (substrate) to pE–CCL2 (product) was estimated by computing the area under the curves (AUC) of substrate and product using Sciex MultiQuant software and the formula: percent conversion = 100% x Product/(Substrate + Product).

### Mass spectrometry

For determination of chemokine susceptibility to DPP4-mediated truncation, hCCL7 (1 μg of chemokine in 20 μl of TRIS buffer, pH 8) was incubated in the presence or absence of 100 nM mDPP4 overnight at 37 °C and were analyzed by liquid chromatography–mass spectrometry. Samples (3 μg) were injected onto a MAbPac reversed-phase liquid chromatography column (Thermo Fisher Scientific, 2.1 mm × 50 mm) heated to 80°C using a Dionex Ultimate 3000 RSLC system. A binary gradient pump was used to deliver solvent A (99.88% water containing 0.1% formic acid and 0.02% trifluoroacetic acid) and solvent B (90% acetonitrile containing 9.88% water plus 0.1% formic acid and 0.02% trifluoroacetic acid) as a gradient of 20% to 65% solvent B over 4.5 min at 300 μL/min. The solvent was step-changed to 90% solvent B over 0.1 min and held at 90% for 6.4 min to clean the column. Finally, the solvent was step-changed to 20% solvent B over 0.1 min and held for 3.9 min for re-equilibration. Eluted samples were analyzed online via electrospray ionization into a Thermo Exactive Plus Extended Mass Range Orbitrap mass spectrometer. Acquired mass spectrometric data were analyzed using Intact Mass software (Protein Metrics).

### *In vitro* CCR2 agonism models

PathHunter eXpress CCR2 CHO-K1 β-Arrestin GPCR Assay (DiscoverX) was performed according to vendor protocol to evaluate the activation of CCR2 by hCCL2 and hCCL7 by measuring β-Arrestin recruitment. Briefly, CHO-K1 CCR2 β-Arrestin cells were thawed and plated in a 96-well plate and incubated at 37℃ for 48 hours to recover from the freeze. For agonist experiments, 10-point, 3-fold serially diluted recombinant human Q–, pE–, or ΔQP–chemokine was added to the cells and incubated for 90 minutes at 37°C. Detection reagent was added and the plate was incubated for 1 hour at room temperature. Luminescence was read on an Envision multimode reader (Perkin Elmer). Percent activity was calculated as follows: 100 x (X - MIN)/(MAX - MIN) where MAX is the luminescent signal from cells treated with 10uM pE–chemokine and MIN is the luminescent signal from cells with no chemokine added. For antagonist experiments, 10-point, 3-fold serially diluted recombinant human ΔQP–CCL2 or ΔQP– CCL7 (antagonist) was added to cells and incubated for 30 minutes at 37℃. Then, either 10 nM pE– CCL2 or 15 nM pE–CCL7 (final concentration, agonist) was added to cells and incubated for an additional 90 minutes at 37°C. Detection reagent was added and the plate was incubated for 1 hour at room temperature. Luminescence was read on an Envision multimode reader. Percent activity was calculated as follows: 100 x (X - MIN)/(MAX - MIN) where MAX is the luminescent signal from cells treated with agonist only and MIN is the luminescent signal from cells with no agonist added.

### Western Blot

THP-1 cells were grown to a density of 2.5×10^5^ viable cells/ml (97.5% viability), pelleted at 250g for 5 min and resuspended in R10 to a density of 3×10^6^ cells/ml. Cells were transferred into a 24-well plate (500µl/well). mCCL7 (Q’–CCL7) and ΔQP’–CCL7 were added to each well (final concentrations were 0 to 50nM). When indicated, pre-stimulation with ΔQP’–CCL7 (50nM, 30min at 37 °C) was done, prior to the addition of Q’–CCL7 (10nM). Upon addition of chemokines, cells were mixed thoroughly and incubated at 37 °C for 10min. The content of each well was transferred into 1.5ml microfuge tubes containing 500µl of cold PBS, and cells pelleted for (1000g, 5min 4 °C). Pellets were lysed in 100µl RIPA buffer on ice for 15min (vortexing was done every 5min). Protein concentration was determined by BCA assay (Pierce #23225). RIPA lysates were prepared for western blotting using Invitrogen Bolt sample loading buffer and reducing agent (per manufacturer’s instructions), denatured for 10min at 95°C with moderate agitation, and loaded into replicate Bolt 4-12% Bis Tris gel (20µg per lane). Gels were run for 2hr at 100v (NEB Color Prestained Protein Standard, Broad Range (11–245 kDa) was used to monitor the run). The gels were transferred onto nitrocellulose membranes using BioRad TransTurbo blot, blocked for 1hr with AdvanBlock Chemi (Advansta #R-03726-E10) and probed with anti-AKT (CellSignalig #4691T) and anti-pAKT-S473 (CellSignaling #9271S, 1:5000 dilution) in AdvanBlock. Blots were incubated overnight at 4°C with gentle rocking. For probing with secondary antibodies, the blots were washed 3x with TBST buffer (5min each) followed by 75min incubation with anti-Rabbit HRP-conjugated antibodies (Jackson Immuno Reserach #711-035-152, 1:5000 dilution). Blots were washed 3x with TBST, 1x with TBS and developed using WesternBright ECL HRP substrate (Advansta #K-12045-D20) on an Azure imager (Azure Biosystems).

### *In vitro* cell migration models

THP-1 cells (1×10^4^) were seeded onto the top chamber of a 96 well transwell plate in RPMI supplemented with 10% FBS and 5mg/ml PMA. 48 hours after, the media was carefully removed from the top chamber, and replaced by HBSS media, supplemented with 0.5% BSA. Chemokines were resuspended in HBSS media supplemented with 0.5% BSA and placed in the bottom chamber of the transwell plate. THP-1 migration was evaluated via microscopy, using a real time measurement of cells that migrated into the lower part of the transwell membrane, with the Incucyte. For horizontal 3D imaging 5×10^5^ cells were pelleted and resuspended in 50µl of center chamber media (10ml RPMI + 2ml FCS + 600µl GlutaMax). Cells were then embedded into a collagen matrix by mixing 20µl of 10xMEM (Gibco), 127µl ddH2O, 3µl NaHCO3, 50µl R10, 50µl Collagen I and 50µl of cell suspension. 6-10µl of cell suspension was loaded into the center chambers of heat/CO2 equilibrated µslide chemotaxis (Ibidi), per manufacturer’s protocol. Slides were incubated for 30min at 37 °C and then side chambers were filled with 60µl of 50nM full Q– CCL7 or ΔQP–CCL7, resuspended in R10. The slides were then imaged using a Nikon TiE inverted microscope (SEE MICROSCOPE SETTINGS). Trajectories were determined using ad the Particle Tracker 2D/3D plugin for ImageJ (using the following settings: radius=20 cutoff=0.5 per/abs=0.500 link=1 displacement=1 dynamics=Brownian). Data was analyzed with R/RStudio and plotted using ggplot2.

### Single cell RNA sequencing

Single cell suspension from WT and *Qpctl^−/−^* LL/2 tumors were prepared as described above. Cells were stained with Live/Dead fixable aqua, and antibodies against CD45.2, CD11c and CD11b before FACS sorting, performed on a BD Aria SORP instrument. Cells were sorted into 1.5 ml tubes (Eppendorf) and counted manually under the microscope. The concentration of cell suspensions was adjusted to 1000 cells/ul. 10000 cells per sample were loaded using the 10x Chromium Single cell 3’ Library, Gel Bead & Multiplex Kit and Chip Kit (10x Genomics, V3.1 barcoding chemistry) according to the manufacturer’s instructions. The subsequent steps to prepare the libraries were performed following the manufacturer’s protocols. Libraries were quantified with Qubit (ThermoFisher dsDNA HS Assay Kit, cat#Q32854) and the average library size was determined using Agilent Tapestation (D1000 screentapes, cat#5067-5582). Purified libraries were analyzed by an Illumina NovaSeq sequencer with 150-bp paired-end reads.

### Bulk tumor-cell line RNA sequencing

Tumor cells were washed once with HBSS. A total of 2×10^6^ cells were homogenized in 140μl of lysis/binding buffer and RNA was extracted using the MagMAX™-96 Total RNA Isolation Kit (AM1830). Quality control of samples was done to determine RNA quantity (using a NanoDrop 8000 (Thermo Scientific)) and quality (using a 2100 Bioanalyzer and 2200 TapeStation (Agilent Technologies)) prior to their processing. Approximately 1ug of total RNA was used as an input material for library preparation using TruSeq RNA Sample Preparation Kit v2 (Illumina). Size of the libraries was confirmed using 2200 TapeStation and High Sensitivity D1K screen tape (Agilent Technologies) and their concentration was determined by qPCR-based method using Library quantification kit (KAPA). The libraries were multiplexed and then sequenced on Illumina HiSeq2500 to generate 30M of single end 50 base pair reads.

### Data representation and statistics

Graphic representation and statistical analysis of the data were done using GraphPad software (v8.0.1). The following statistical tests were used: Unparametric Mann-Whitney test, Mixed-effects ANOVA paired test with Geisser-Greenhouse correction; and two-way ANOVA with Geisser-Greenhouse correction, as indicated in the figure legends.

### Single Cell RNA-seq data processing and analysis

Single-cell RNA-seq data was processed with cellranger count (CellRanger 3.0.2 from 10x Genomics) using a custom reference package based on mouse reference genome GRCm38 and GENCODE ^53^ gene models. Data analysis was done in R 3.6.2 and the Seurat package version 3.1.5 ^54^. Only high-quality cells were retained for the posterior analysis, more concretely, we kept the cells with more than 300 hundred genes detected, more than 1000 unique molecule identifiers (UMIs) and less than 5% mitochondrial read. This filtering step removed most of the doublets and empty cells. Using the talus plot ^55^ we determined that 60 principal components were enough to capture the signal in data. With these retained components, we computed a UMAP embedding and the neighbors for the clustering. Several clustering resolutions were calculated and a directed tree was constructed reflecting the hierarchical relationships of the new clusters upon increasing the resolution ^56^. A resolution of 0.1 was considered optimal and the main clusters were identified by means of expression markers of known biology. Markers for each cluster were identified using the Seurat::FindAllMarkers() method with default parameters, comparing all cells in a particular cluster to the rest of cells in the dataset and accessing significantly differential gene expression using Wilcoxon’s rank sum test and Benjamini Hochberg correction for multiple testing. To focus on Mono/Mac/DCs cells, clusters that were enriched with markers of other immune or stromal cells were excluded from the analysis: B cells (*Cd79a/b*), stroma (*Dkk2*, *Dcn*), Lymphocytes (*Gzma/b*), neutrophils (*S100a8/9*). A resolution of 0.6 was chosen to identify the cell clusters. The identity of individual clusters was determined by calculating scores of previously published gene sets in each of our identified clusters using the AddModuleScore function in Seurat with default parameters. Gene sets for phagocytosis, classically activated macrophages (M1), alternatively activated macrophages (M2) and angiogenesis ^11^ as well as antigen presentation (GO-term) were used. The similarity score of the Ki67_MAC cluster was accessed by mapping it back to the other clusters using the methodology of SingleR^57^ with the remaining clusters as the target reference. The analysis of cluster similarity between loss of QPCTL and CSF1R treatment was done using macrophage gene sets published by Zhang and colleagues ^17^.

## ACKNOWLEDGEMENTS

The authors would like to acknowledge the following people, for technical and intellectual support: Yan Lu, Jiansheng Wu, Jian Payandeh, Isabelle Lehoux and protein structure/purification teams; Benjamin Haley, Jennie Lill, Wendy Sandoval and Peter Liu; Soren Warming, Udi Segal, and animal care-takers; Yajun Chestnut, Manmeet Singh and Alan Gutierrez; Pathology, necropsy and antibody engineering/production teams; Allison Cordrey and cell culture team. We also thank Wenxian Fu for careful reading of the manuscript. Illustrations were generated by the authors, or adapted from Servier medical art (France) or Biorender.

## AUTHOR CONTRIBUTIONS

Conceptualization: R.B.d.S. and M.L.A.; Investigation and Methodology Development: R.B.d.S., R.L., S.W., J.O., V.J., Y.W., W.P., C.Ev., J.N., D.A., J.Z., Z.M., J.M.S. and M.M.; Data Analysis: R.B.d.S., R.L., X.P., J.O., C.Ev., J.N. and J.Z.; Resources: C.L., Y-C.H., J.T.K., I.H., T.P. and M.R.G.; Supervision: R.B.d.S., M.L.A., C.Ei., I.M. J.M.S. and M.M.; Writing – Original Draft: R.B.d.S.; Writing – Review & Editing Preparation: M.L.A., C.Ei., S.R. and I.M., with edits from all authors.

## DATA AND MATERIAL AVAILABILITY STATEMENT

The data that support the findings of this study are available from the corresponding authors upon reasonable request. Material requests should be done to the correspondent authors.

