## Supplementary data for "Loss of the intracellular enzyme QPCTL limits chemokine function and reshapes myeloid infiltration to augment tumor immunity"

### EXTENDED DATA

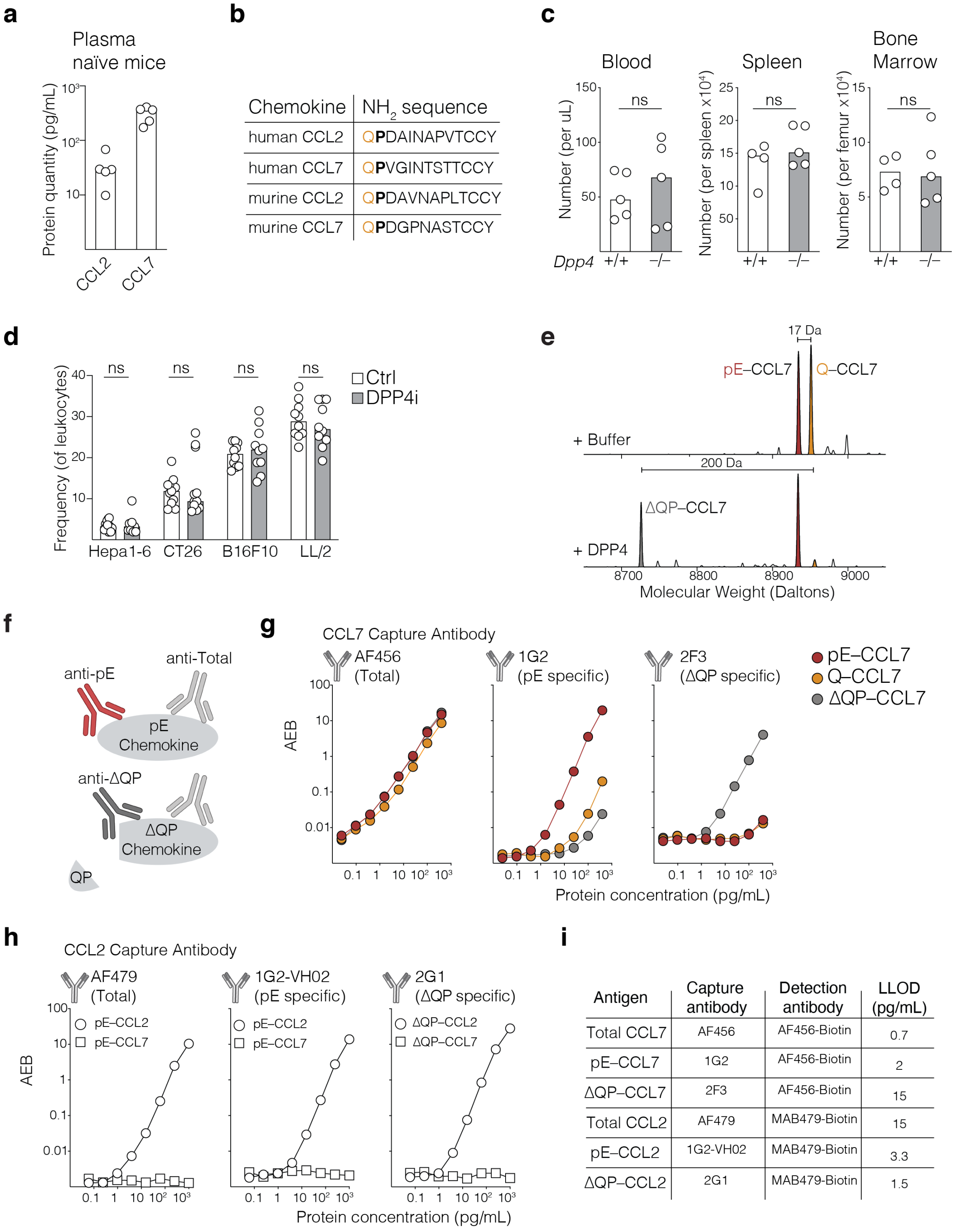


**Extended Data Fig. 1 - Monocyte migration is not modulated by DPP4**

**a**, Quantification of mouse (m)CCL2 and mCCL7 in plasma of naïve WT mice (n = 5 mice per group).

**b**, $NH_2$-terminal amino acid sequence of the human and mouse monocyte chemoattractants CCL2 and CCL7. The presence of a glutamine (Q) in the first position is highlighted in orange and the proline (P) in the second position is highlighted in bold.

**c**, Quantification of monocytes ($CD11b^+Ly6G^-SiglecF^-Ly6C^+$) in blood (n = 5), spleen (n = 4 or 5) and bone marrow (n = 4 or 5) of naïve WT littermate (*$Dpp4^{+/+}$*) *and $Dpp4^{-/-}$* mice.

**d**, Quantification of monocytes in tumors implanted in WT mice treated with control (ctrl) chow or chow containing DPP4i (n= 20 (Hepa1-6), 22 (CT26), 20 (B16f10) or 21 (LL/2) mice).

**e**, Recombinant human (h)CCL7 was incubated in the absence (upper lane) or presence of recombinant hDPP4 (second lane). Samples were analyzed by mass spectrometry and molecular weight profiles of resulting species are shown in Daltons (Da).

**f**, Schematic representation of antibodies that detect post-translational modifications (PTMs) in mouse chemokines: anti-pE recognizes N-terminal cyclization and anti-delta(Δ)QP is specific for the N-terminal truncated chemokine. Anti-total antibody recognizes the chemokine independently of its N-terminal modifications.

**g**, Cross-reactivity profile of anti-mCCL7 capture antibodies against the indicated concentrations of each mCCL7 N-terminal forms. Capture of mCCL7 was done with either AF456, an antibody that recognizes mCCL7 independently of its N-terminal modifications (total) or with 1G2, which binds preferentially to the N-terminal pE-mCCL7 (pE-specific) or with 2F3, which binds preferentially to the N-terminal truncated CCL7 (delta(Δ)QP specific). Detection was done with biotinylated anti-mCCL7 AF456, using the SIMOA technology. AEB, average enzyme per bead.

**h**, Cross-reactivity profile of mCCL2 capture antibodies against the indicated concentrations of mCCL7 and mCCL2 forms. Detection was done with biotinylated anti-mCCL2 MAB479, using the SIMOA technology.

**i**, Description of capture and detection antibodies used to analyze mCCL2 and mCCL7 PTMs. Lower limit of detection (LLOD) values are shown.

Bars are medians and symbols individual mice. Data shown is a representative experiment (**a**,**c**,**g**,**h**) or pooled from 2 experiments (**d**). All experiments were repeated independently (⩾2 times). ns, not significant. P values are from nonparametric Mann-Whitney test.

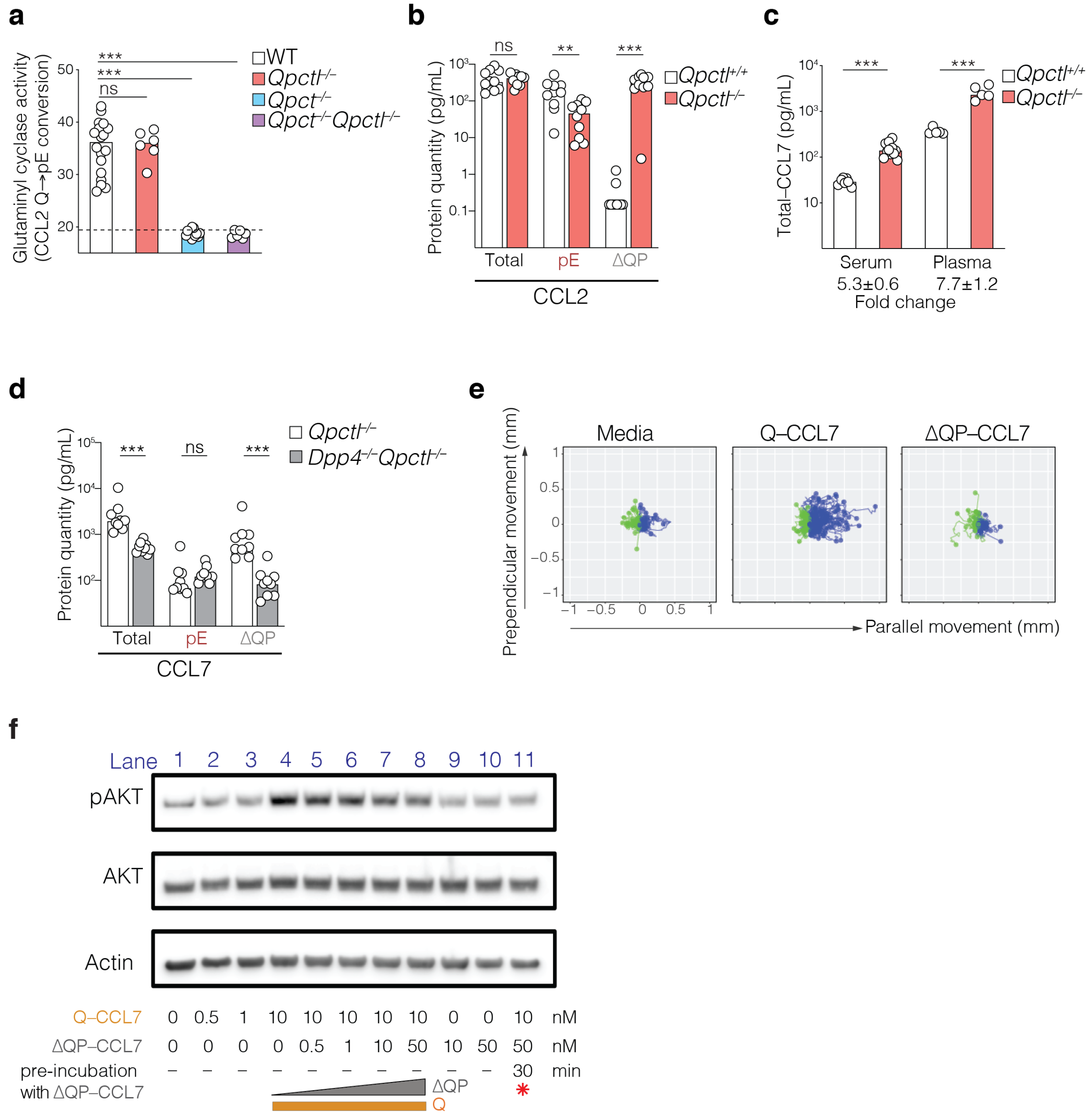


**Extended Data Fig. 2 – *In vivo* regulation of mCCL7 PTMs by QPCTL and DPP4**

**a**, Glutaminyl cyclase activity was determined in serum from the indicated mouse genotypes, by measuring the conversion of spiked hCCL2 N-terminal Glutamine (Q) to pyroglutamate (pE) by mass spectrometry (n= 18, 6, 12 and 6 mice, respectively for each depicted genotype).

**b,** Quantification of CCL2 PTMs in plasma of *Qpctl*–/– and *Qpctl*+/+ littermate mice bearing *Qpctl*–/– and *Qpctl*+/+ LL/2 tumors, respectively.

**c**, Quantification of total–CCL7 in serum (n = 7 or 11 mice per genotype) and plasma (n = 5 mice per genotype) from *Qpctl*$^{+/+}$ and *Qpctl*$^{-/-}$ littermate mice.

**d**, Quantification of mCCL7 PTMs in plasma of naïve *Qpctl*$^{-/-}$ and littermate *Dpp4*$^{-/-}$*Qpctl*$^{-/-}$ mice (n = 9 mice per genotype).

**e**, Migration of THP-1 cells to the indicated hCCL7 forms placed on the right side of a u-migration chamber. Cellular trajectories that moved towards the chemokine are represented in blue while greens trajectories are cells that moved away from the chemokine.

**f**, Akt phosphorylation (pAkt) in THP-1 cells was measured by Western blot, after incubation with media (lane 1); Q–CCL7 (lanes 2 to 4) or ΔQP–CCL7 (lanes 9 and 10). Antagonist activity of ΔQP–CCL7 was evaluated by measuring pAkt following incubation with 10nM of Q–CCL7 and the indicated doses of ΔQP–CCL7 (lanes 5 to 8). Q–CCL7 is not able to induce pAkt in THP-1 cells if the cells were pre-incubated with ΔQP–CCL7 (lane 11, red asterisk).

Bars are medians and symbols individual mice. Data shown are representative experiments. All experiments were performed independently (⩾2 times). ns, not significant; ** $p \leqslant 0.01$; *** $p \leqslant 0.001$. P values are from nonparametric Mann-Whitney test.

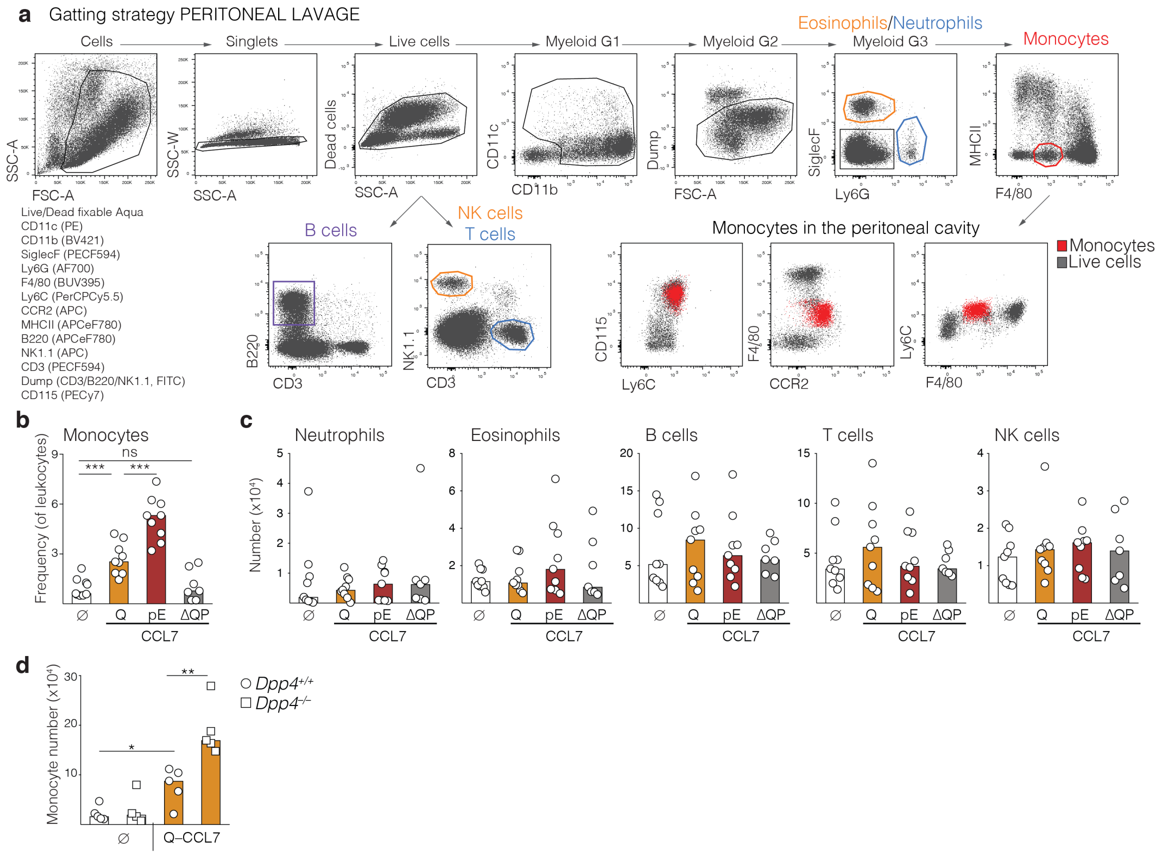


**Extended Data Fig. 3 – Model of mCCL7-mediated migration of leukocytes to the peritoneal cavity**

**a**,**b**,**c**, WT mice were injected with PBS (ø) or with recombinant mCCL7 forms in the peritoneal cavity. **a**, Gating strategy for the identification of peritoneal leukocytes. **b,** Frequency of peritoneal monocytes and **c**, number of neutrophils, eosinophils, B cells, T cells and NK cells is shown (n = 9 or 7 (ΔQP) mice per group).

**d**, Mouse Q–CCL7 was injected in the peritoneal cavity of WT littermate (*Dpp4*$^{+/+}$*) or Dpp4*$^{-/-}$ mice. The number of peritoneal monocytes was quantified by flow cytometry (n = 5 or 4 (*Dpp4*$^{-/-}$ ø) mice per group.

Bars are medians and symbols individual mice. Data shown are representative experiments (**d**) or pooled from 2 experiments (**b** and **c**). All experiments were repeated (⩾2 times). ns, not significant; * $p \leqslant 0.05$ ** $p \leqslant 0.01$; *** $p \leqslant 0.001$. P values are from nonparametric Mann-Whitney test.

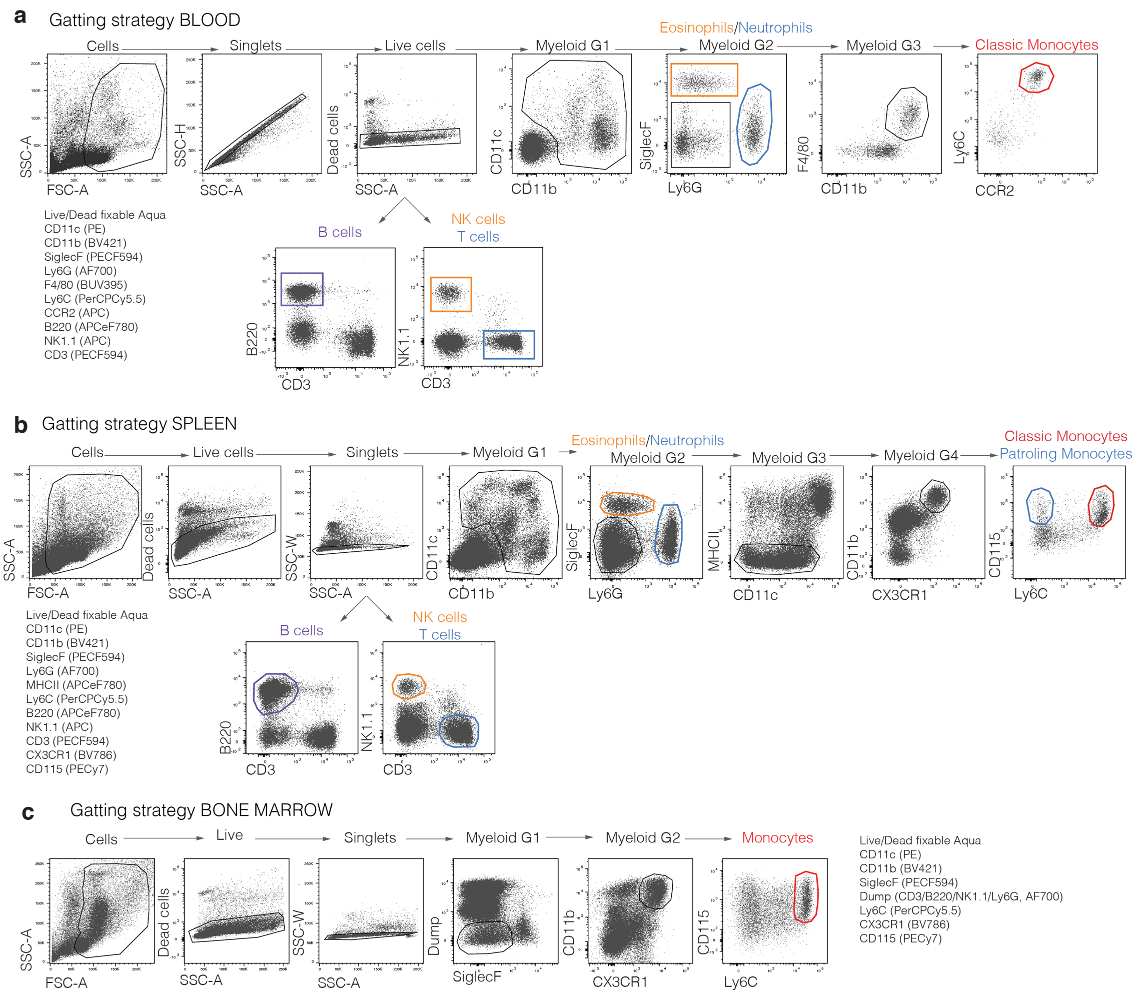


**Extended Data Fig. 4 – Gating strategy for identification of leukocytes subsets in mouse tissues**

**a**,**b**,**c**, Gating strategy for identification of leukocytes subsets in mouse **a**, blood, **b**, spleen and **c**, bone marrow.

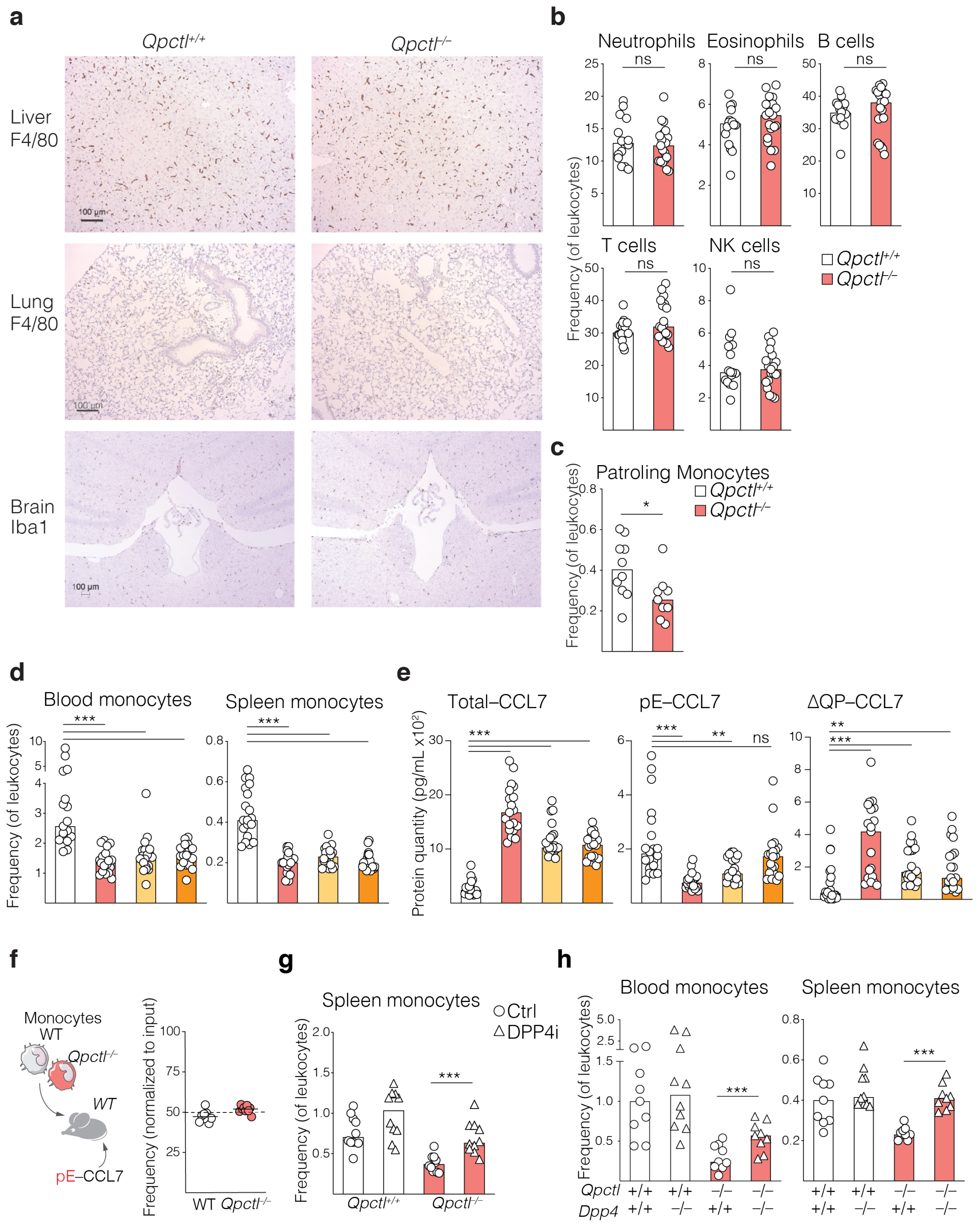


**Extended Data Fig. 5 – Loss of *Qpctl* in both hematopoietic or stromal compartments impairs monocyte homeostasis in mice but it does not impact tissue-resident macrophages, nor lymphocyte populations.**

**a**, Histology analysis of liver, lung and brain sections from naive WT littermate (*Qpctl*$^{+/+}$) and *Qpctl*$^{-/-}$ mice. Identification of tissue macrophages was done by immuno-staining of F4/80 (liver and lung) or Iba1 (brain) expressing cells. Scale bars are 100uM.

**b**, Frequency of leukocytes in blood collected from *Qpctl*$^{+/+}$ and *Qpctl*$^{-/-}$ mice was determined by flow cytometry (n = 16 or 19 (*Qpctl*$^{-/-}$) mice per group).

**c**, Frequency of patrolling monocytes (CD11b$^{+}$CD115$^{+}$Ly6C$^{-}$CX3CR1$^{+}$) among splenic leukocytes from *Qpctl*$^{+/+}$ and *Qpctl*$^{-/-}$ mice is shown (n = 10 or 9 (*Qpctl*$^{-/-}$) mice per group).

**d**,**e**, Bone marrow chimeras were generated by reconstituting lethally irradiated WT or *Qpctl*$^{-/-}$ mice with hematopoietic progenitors from WT or *Qpctl*$^{-/-}$ mice. **d**, Quantification of monocytes in the blood and spleen of chimeric mice. **e**, Quantification of CCL7 PTMs in the plasma of chimeric mice (n = 19 (*Qpctl*$^{-/-}$ into *Qpctl*$^{-/-}$) or 20 mice per group).

**f**, WT (CD45.1) and *Qpctl*$^{-/-}$ (CD45.2) monocytes were co-transferred into WT (CD45.1/CD45.2) hosts. Migration of transferred monocytes into the peritoneal cavity following intraperitoneal injection of pE–CCL7 was assessed by flow cytometry. Frequency of peritoneal WT (CD45.1) and *Qpctl*$^{-/-}$ (CD45.2) monocytes was normalized to the ratio of WT and *Qpctl*$^{-/-}$ monocytes transferred (n = 9 mice per group).

**g**, WT littermate (*Qpctl*$^{+/+}$) and *Qpctl*$^{-/-}$ mice were treated with control (Ctrl) chow or chow containing DPP4i. Frequency of splenic monocytes after 4 weeks of treatment is shown, n = 10 mice per group.

**h**, Quantification of monocyte frequency in blood and spleen of naïve littermate WT, *Dpp4*$^{-/-}$, *Qpctl*$^{-/-}$ and *Dpp4*$^{-/-}$*Qpctl*$^{-/-}$ double mutant mice (n = 10 (*Dpp4*$^{-/-}$) or 9 mice per group).

Bars are medians and symbols individual mice. Data shown are representative experiments (**a**, **f**), or pooled from 2-3 experiments (**b**,**c**,**d**,**e,g,h**). All experiments were done at least 2 times, with the exception of histology in mouse tissues that was done once (n = 5 mice per group). ns, not significant; * $p \leqslant 0.05$. ** $p \leqslant 0.01$, *** $p \leqslant 0.001$. P values are from nonparametric Mann-Whitney test.

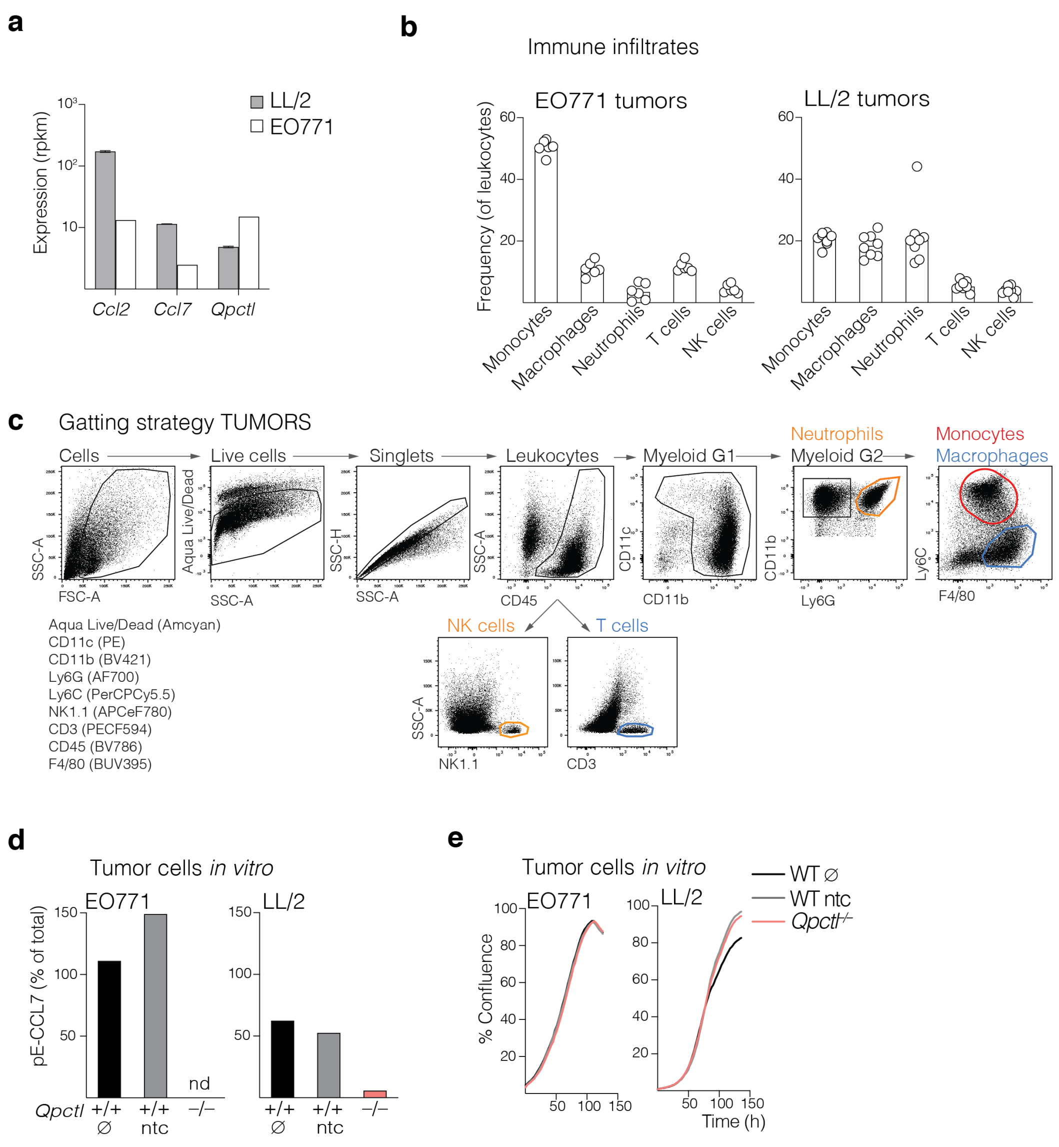


**Extended Data Fig. 6 – Characterization of EO771 and LL/2 mouse tumor models**

**a**, Quantification of *Ccl2*, *Ccl7* and *Qpctl* RNA expression in LL/2 and EO771 cells.

**b**, WT mice were inoculated with EO771 (left graph, n = 6 mice per group) or with LL/2 (right graph, n = 8 mice per group). Frequency of tumor associated leukocytes was quantified by flow cytometry, 14 days after inoculation.

**c**, Gating strategy for the identification of leukocytes in tumors.

**d**,**e**, CRISPR / Cas9 edited *Qpctl*$^{-/-}$ mutant LL/2 and EO771 tumor lines were generated. **d**, Quantification of pE–CCL7 was done in the supernatants from the cells and plotted as percentage of total–CCL7, measured in the same samples. **e**, *In vitro* growth curves of tumor cell lines were determined using the Incucyte technology. ∅ - parental cell line; ntc - non-targeted control.

Bars are medians and symbols individual mice. Data shown are representative experiments. All experiments were repeated independently (⩾3 times), except quantification of RNA expression (**a**), which was done once.

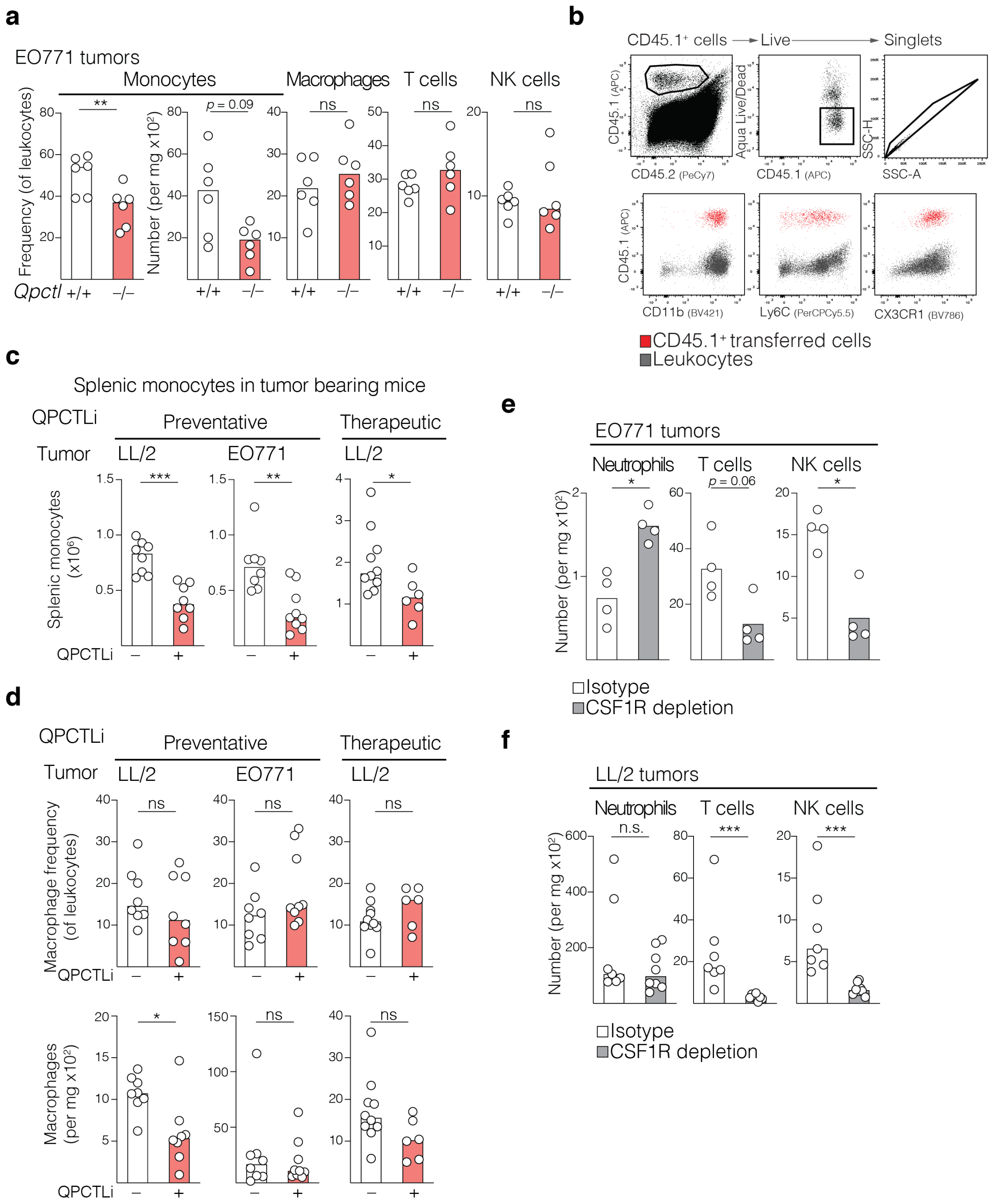


**Extended Data Fig. 7 – Modulation of QPCTL impairs monocyte migration into tumors**

**a**, *Qpctl*$^{+/+}$ and *Qpctl*$^{-/-}$ littermate mice were inoculated with *Qpctl*$^{+/+}$ or *Qpctl*$^{-/-}$ EO771 tumor cells, respectively. The frequency of tumor infiltrating Ly6C$^{+}$ monocytic cells and the number of Ly6C$^{+}$ monocytes, macrophages, T cells and NK cells is depicted (n = 6 mice per group).

**b**, Gating strategy to identify tumor infiltrating WT CD45.1$^{+}$ monocytes transferred into LL/2 tumor bearing mice.

**c,d**, WT mice received ctrl vehicle or QPCTLi for 4 days before LL/2 or EO771 tumor inoculation (preventative) or at day 7 after LL/2 inoculation (therapeutic). On day 14 after tumor cell inoculation **c**, the number of splenic monocytes and **d**, the number of tumor-infiltrating macrophages were determined by flow cytometry (n = 8 (LL/2 preventative); n = 8 or 9 (QPCTLi EO771 preventative) and n = 10 or 6 (QPCTLi LL/2 therapeutic)).

**e**,**f**, WT mice treated with ctrl isotype or anti-CSF1R antibody were inoculated with **e**, EO771 cells or **f**, LL/2 cells. Quantification of leukocytes in tumors excised **e**, 14 days or **f**,19 days after tumor cell inoculation is shown (n = 4 (LL/2) or n = 7 or 8 mice (EO771, anti-CSF1R) per group).

Bars are medians and symbols individual mice. Data shown are representative experiments. All experiments were repeated independently (⩾2 times). *$p \leqslant 0.05$, **$p \leqslant 0.01$, ***$p \leqslant 0.001$. P values are from nonparametric Mann-Whitney test.

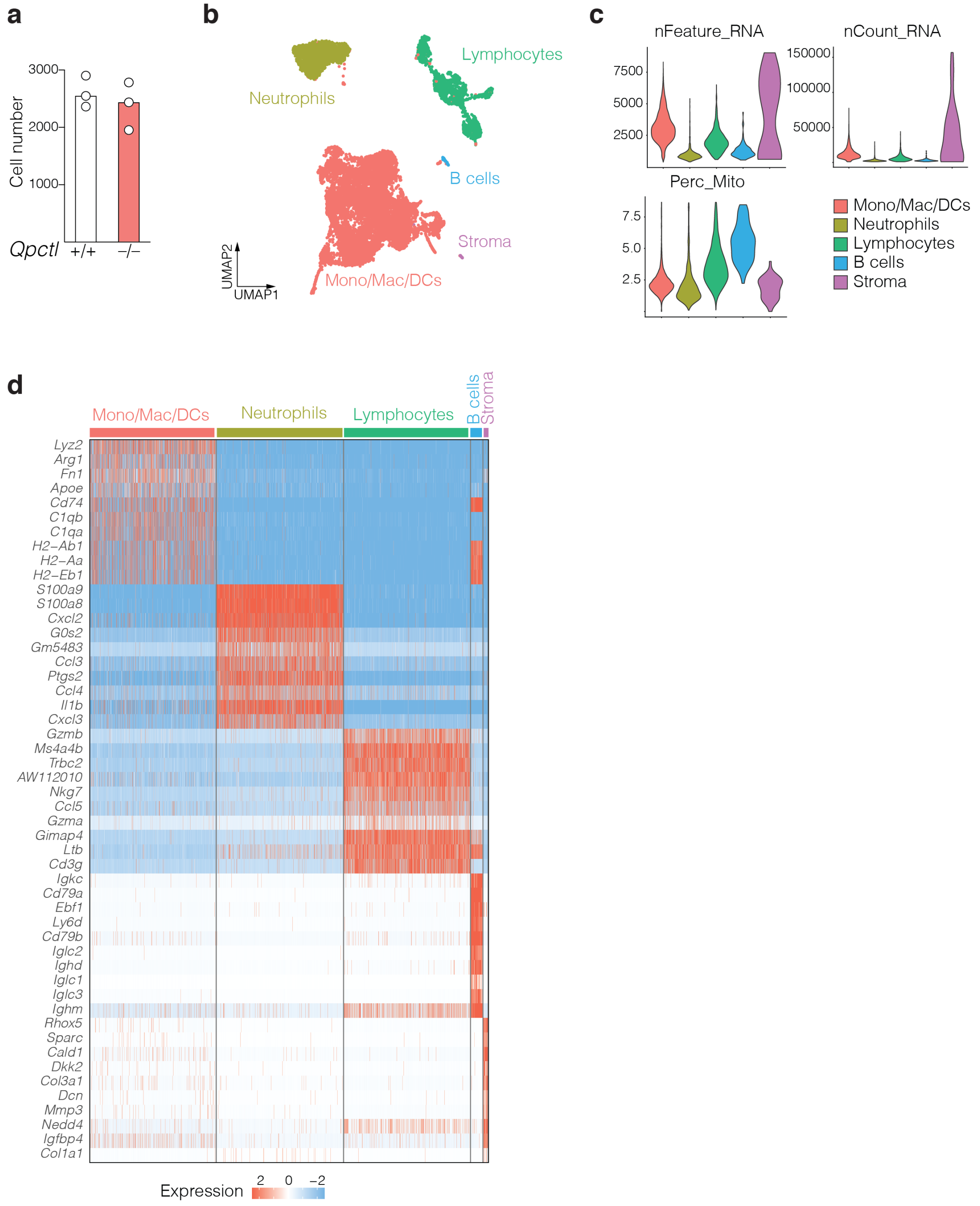


**Extended Data Fig. 8 – Analysis of immune infiltrates by single cell RNAseq**

**a**,**b**,**c**,**d**, *Qpctl*$^{-/-}$ and *Qpctl*$^{+/+}$ littermate mice were inoculated with *Qpctl*$^{-/-}$ or *Qpctl*$^{+/+}$ LL/2 tumors, respectively. On day 14 after tumor inoculation, tumors were excised and processed for single cell RNAseq. **a**, Bar graph representing the number of cells recovered per sample. **b**, UMAP plots of tumor immune infiltrates, from merged samples (n = 6, 3 samples per genotype). **c**, Violin plots representing quality control measurements. **d**, Heatmap showing normalized expression of top 10 expressed genes in each cluster. Bars are medians and symbols individual mice.

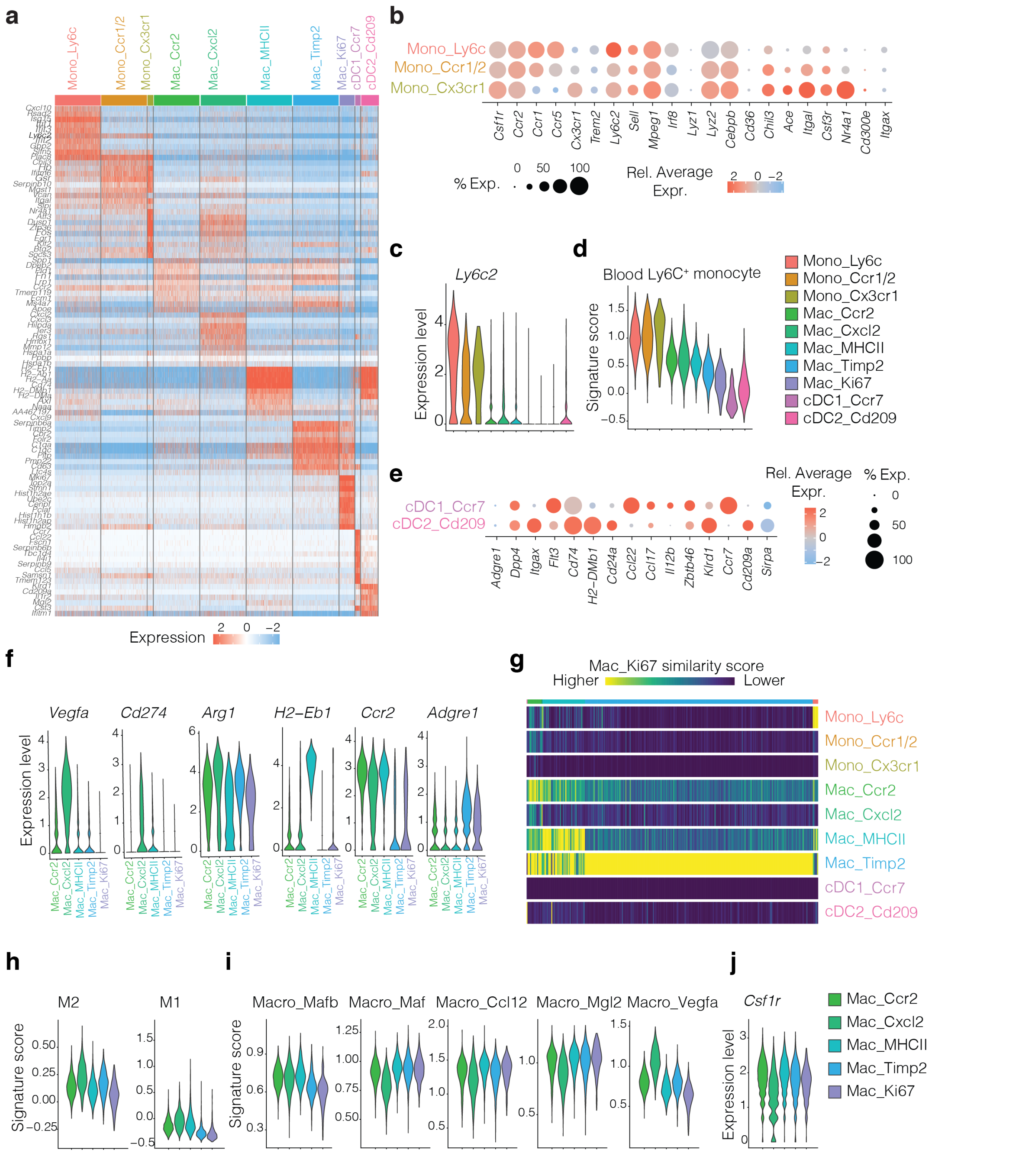


**Extended Data Fig. 9 – Loss of *Qpctl* remodels the tumor myeloid compartments**

**a**,**b**,**c**,**d**,**e**,**f**,**g**,**h**,**i**,**j** *Qpctl*$^{-/-}$ and *Qpctl*$^{+/+}$ littermate mice were inoculated with *Qpctl*$^{-/-}$ or *Qpctl*$^{+/+}$ LL/2 tumors, respectively. On day 14 after tumor inoculation, tumors were excised and processed for single cell RNAseq. **a**, Heatmap showing normalized expression of top 10 expressed genes in each Mono/Mac/DCs cluster. **b**, Dot plots of selected markers in monocyte populations. Dot size indicates proportion of cells in each cluster expressing a gene, color shading indicates the relative level of gene expression. **c**, Violin plot representing *Ly6c2* expression across Mono/Mac/DCs clusters. **d**, Violin plot representing the expression of a blood Ly6C$^{+}$ monocyte signature across Mono/Mac/DCs clusters. **e**, Dot plots of selected markers in dendritic cell populations. Dot size indicates proportion of cells in each cluster expressing a gene, color shading indicates the relative levels of gene expression. **f**, Violin plots representing the expression of indicated genes across macrophage clusters. **g**, The similarity score between the Mac_Ki67 cluster and the other Mono/Mac/DC clusters was calculated using SingleR automatic annotation. **h**,**i**, Violin plots representing the expression of indicated gene signatures across macrophage clusters. **j**, Violin plot representing *Csf1r* expression across macrophage clusters.

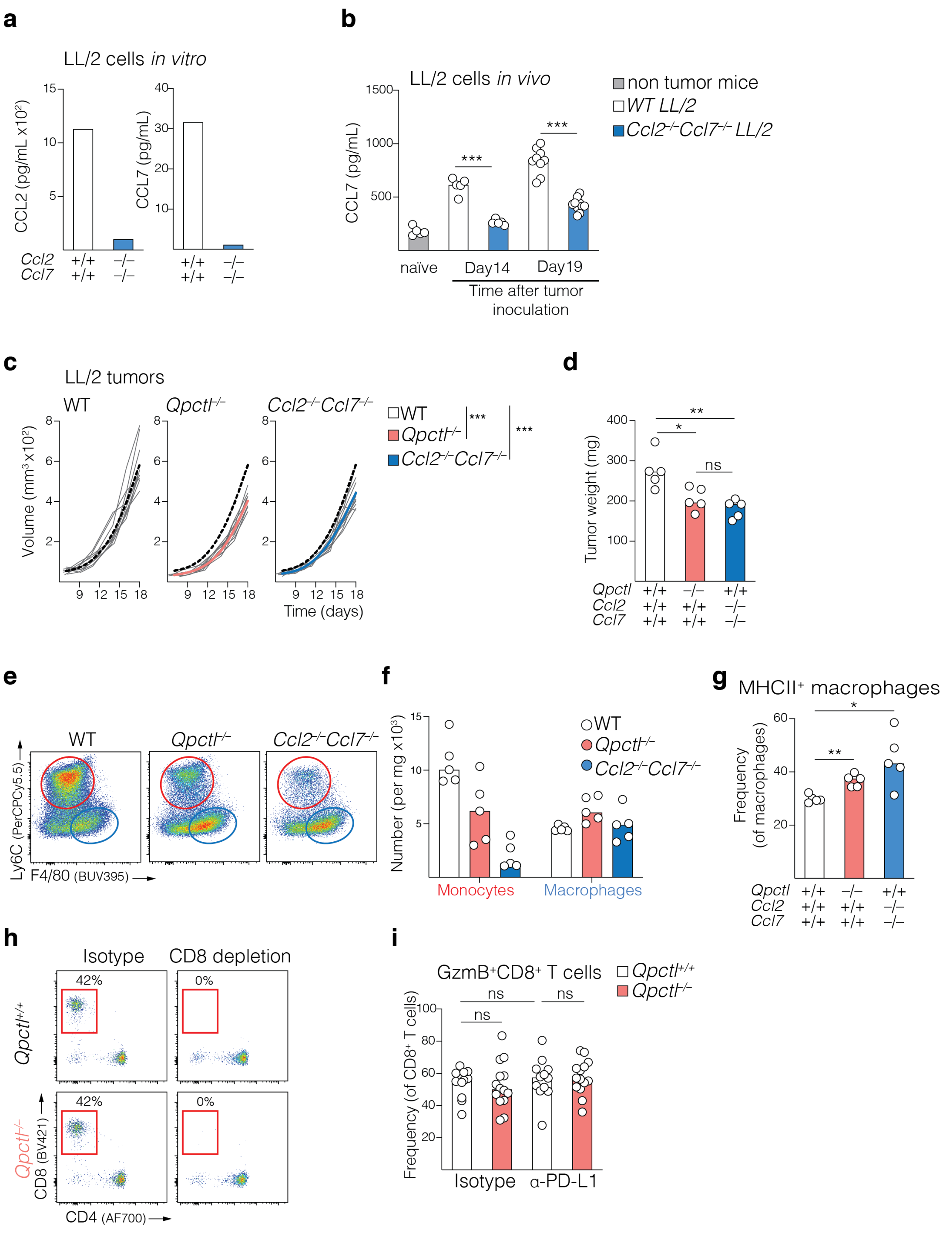


**Extended Data Fig. 10 – Loss of *Ccl2* and *Ccl7* mimics loss of *Qpctl* in LL/2 tumors**

**a**,**b**, CRISPR / Cas9 edited *Ccl2*$^{-/-}$*Ccl7*$^{-/-}$ double mutant LL/2 tumor cells were generated. **a**, Quantification of total–CCL2 and total–CCL7 was done in the supernatants from the cells. **b**, WT mice were inoculated with WT or *Ccl2*$^{-/-}$*Ccl7*$^{-/-}$ LL/2 cells. CCL7 was quantified in the plasma samples of naïve and tumor bearing mice 14 or 19 days after tumor cell inoculation (n= 5 and 10 mice per group, respectively per time point).

**c**,**d**, WT mice were inoculated with WT, *Qpctl*$^{-/-}$ or *Ccl2*$^{-/-}$*Ccl7*$^{-/-}$ LL/2 cells. **c**, Tumor growth was measured over time (n = 10 mice per group). Gray lines represent individual mice and overlay of fitting spline cubic curves for each group is shown. Black dotted lines represent spline curves from the WT group. **d**, Weight of tumors excised at day 14 is shown (n = 5 mice per group).

**e**,**f**,**g**, WT mice were inoculated as described in (**c**). Tumors were analyzed by flow cytometry 14 days after implantation. **e**, Representative flow cytometry plots (gated on live, singlets, $CD45^+CD11b^+Ly6G^-$) highlighting $Ly6C^+$ monocytic cells (red circle) and $F4/80^{high}$ macrophages (blue circle). **f**, Quantification of $Ly6C^+$ monocytic cells and macrophages is shown (n= 5 mice per group). **g**, Frequency of $MHCII^+$ macrophages is shown (n = 5 mice per group).

**h**, *Qpctl*$^{+/+}$ and *Qpctl*$^{-/-}$ EO771 bearing mice were treated with ctrl isotype or anti-CD8 depletion antibody. Representative flow cytometry plots (gated on live, single, $CD3^+$) of the T cell composition in blood, assessed 2 days after antibody injection are shown.

**i**, *Qpctl*$^{+/+}$ and *Qpctl*$^{-/-}$ littermate mice were inoculated with *Qpctl*$^{+/+}$ or *Qpctl*$^{-/-}$ EO771 tumor cells, respectively. On day 9 after tumor cell inoculation, mice were injected with control isotype or anti-PD-L1 blocking antibody. Tumors were excited 7 days after initiation of treatment and analyzed by flow cytometry. Frequency of granzyme B-expressing intra-tumoral $CD8^+$ T cells is shown (n = 12 (WT) or 14 (*Qpctl*$^{-/-}$) mice).

Bars are medians and symbols individual mice. Data shown are representative experiments or pooled from 2 (**i**). All experiments were repeated independently (⩾2 times). ns, not significant $^*p \leqslant 0.05$, $^{**}p \leqslant 0.01$, $^{***}p \leqslant 0.001$. P values are from nonparametric Mann-Whitney test (**b**,**d**,**g**,**i**) or Two-way ANOVA test (**c**).

**Supplementary Table 1.** Gene signatures used to score blood monocytes, M1, M2, Angiogenesis, Phagocytosis and Antigen Presentation (GO:0019882).

| **Monocytes** | **M1** | **M2** | **Angiogenesis** | **Phagocytosis** | **Antigen Presentation** |
|---|---|---|---|---|---|
| *Ly6c2* | *Il23a* | *Il4r* | *Ccn2* | *Mrc1* | *Abcb9* |
| *Chi3l3* | *Tnf* | *Ccl4* | *Ccn1* | *Cd163* | *Ap3b1* |
| *Lyz2* | *Cxcl9* | *Ccl13* | *Cd44* | *Mertk* | *Atg5* |
| *F13a1* | *Cxcl10* | *Ccl20* | *Cxcr4* | *C1qb* | *Azgp1* |
| *Hp* | *Cxcl11* | *Ccl17* | *E3f3* | | *B2m* |
| *Ly6c1* | *Cd86* | *Ccl18* | *Edn1* | | *Bag6* |
| *Emb* | *Il1a* | *Ccl22* | *Ezh2* | | *Calr* |
| *Gm11428* | *Il1b* | *Ccl24* | *Fgf18* | | *Ccl19* |
| *C3* | *Il6* | *Lyve1* | *Fgfr1* | | *Ccl21a* |
| *Fn1* | *Ccl5* | *Vegfa* | *Fyn* | | *Ccr7* |
| *Ccl6* | *Irf5* | *Vegfb* | *Hey1* | | *Cd1d1* |
| *Tmsb10* | *Irf1* | *Vegfc* | *Itgav* | | *Cd68* |
| *Mgst1* | *Cd40* | *Vegfd* | *Jag1* | | *Cd74* |
| *AC113316,1* | *Ido* | *Egf* | *Jag2* | | *Clec4a2* |
| *Ccl9* | *Kynu* | *Ctsa* | *Mmp9* | | *Clec4b2* |
| *Lyz1* | *Ccr7* | *Ctsb* | *Notch1* | | *Ctse* |
| *Npc2* | | *Ctsc* | *Pdgfa* | | *Ctsl* |
| *Ms4a4c* | | *Ctsd* | *Ptk2* | | *Ctss* |
| *Mpeg1* | | *Tgfb1* | *Spp1* | | *Erap1* |
| *S100a6* | | *Tgfb2* | *Stc1* | | *Ext1* |
| *Gm10925* | | *Tgfb3* | *Tnfaip6* | | *Fcer1g* |
| *Ccr2* | | *mmp14* | *Tymp* | | *Fcgr1* |
| *Sell* | | *Mmp19* | *Vav2* | | *Fcgr2b* |
| *Ifi27l2a* | | *Mmp9* | *Vcan* | | *Fcgr3* |
| *Prdx5* | | *Clec7a* | *Vegfa* | | *Fgl2* |
| | | *Wnt7b* | | | *Flt3* |
| | | *Fasl* | | | *Gba* |
| | | *Tnfsf12* | | | *Gm7030* |
| | | *Tnfsf8* | | | *Gm8909* |
| | | *Cd276* | | | *Gm11127* |

| | | | | | |
|---|---|---|---|---|---|
| | | *Vtcn1* | | | *H2-Aa* |
| | | *Msr1* | | | *H2-Ab1* |
| | | *Irf4* | | | *H2-D1* |
| | | | | | *H2-DMa* |
| | | | | | *H2-DMb1* |
| | | | | | *H2-DMb2* |
| | | | | | *H2-Ea* |
| | | | | | *H2-Eb1* |
| | | | | | *H2-K1* |
| | | | | | *H2-L* |
| | | | | | *H2-M1* |
| | | | | | *H2-M2* |
| | | | | | *H2-M3* |
| | | | | | *H2-M5* |
| | | | | | *H2-M9* |
| | | | | | *H2-M10.1* |
| | | | | | *H2-M10.2* |
| | | | | | *H2-M10.3* |
| | | | | | *H2-M10.4* |
| | | | | | *H2-M10.5* |
| | | | | | *H2-M10.6* |
| | | | | | *H2-M11* |
| | | | | | *H2-Oa* |
| | | | | | *H2-Ob* |
| | | | | | *H2-Q1* |
| | | | | | *H2-Q2* |
| | | | | | *H2-Q4* |
| | | | | | *H2-Q6* |
| | | | | | *H2-Q7* |
| | | | | | *H2-Q8* |
| | | | | | *H2-Q9* |
| | | | | | *H2-Q10* |
| | | | | | *H2-T3* |
| | | | | | *H2-T22* |

| | | | | | |
|---|---|---|---|---|---|
| | | | | | *H2-T23* |
| | | | | | *H2-T24* |
| | | | | | *Hfe* |
| | | | | | *Icam1* |
| | | | | | *Ide* |
| | | | | | *Ifi30* |
| | | | | | *Ifng* |
| | | | | | *Ighg2a* |
| | | | | | *Ighm* |
| | | | | | *Kdm5d* |
| | | | | | *Marchf1* |
| | | | | | *Marchf8* |
| | | | | | *Mfsd6* |
| | | | | | *Mr1* |
| | | | | | *Nod1* |
| | | | | | *Nod2* |
| | | | | | *Pikfyve* |
| | | | | | *Psap* |
| | | | | | *Psmb8* |
| | | | | | *Psmb9* |
| | | | | | *Psme1* |
| | | | | | *Psme2* |
| | | | | | *Ptpn22* |
| | | | | | *Pycard* |
| | | | | | *Rab3b* |
| | | | | | *Rab3c* |
| | | | | | *Rab4a* |
| | | | | | *Rab5b* |
| | | | | | *Rab6a* |
| | | | | | *Rab8b* |
| | | | | | *Rab10* |
| | | | | | *Rab27a* |
| | | | | | *Rab32* |
| | | | | | *Rab33a* |

| | | | | | |
|---|---|---|---|---|---|
| | | | | | *Rab34* |
| | | | | | *Rab35* |
| | | | | | *Relb* |
| | | | | | *Rftn1* |
| | | | | | *Slc11a1* |
| | | | | | *Tap1* |
| | | | | | *Tap2* |
| | | | | | *Tapbp* |
| | | | | | *Tapbpl* |
| | | | | | *Thbs1* |
| | | | | | *Traf6* |
| | | | | | *Trem2* |
| | | | | | *Treml4* |
| | | | | | *Trex1* |
| | | | | | *Unc93b1* |
| | | | | | *Was* |
| | | | | | *Washc1* |
| | | | | | *Wdfy4* |
| | | | | | *Ythdf1* |
| | | | | | *Psmb9* |
